# High-throughput LacZ/CPRG screen identifies novel potential antibiotics targeting Gram-negative bacterial envelopes to combat resistance

**DOI:** 10.1101/2025.04.09.648071

**Authors:** Fardin Ghobakhlou, Larbi Mokhtari, Tom A. Pfeifer, Catherine Paradis-Bleau

## Abstract

**Background:** Bacterial resistance, exacerbated by multidrug-resistant Gram-negative (GN) pathogens, poses a public health threat due to their impermeable envelopes, which block many antibiotics.

**Objectives:** We aimed to develop a high-throughput screening (HTS) method to identify small molecules targeting GN bacterial envelopes and assess their antibacterial potential.

**Methods:** Envelope disruption in *Escherichia coli* and *Pseudomonas aeruginosa* was assessed using a *β*-galactosidase (LacZ)/CPRG reporter assay in LB at 37°C. The assay was validated through screening the LOPAC^1280^ and KD2^4761^ compound libraries. Concentration–response relationships, permeabilisation constants (K_50_), co-permeabilisation assays, minimal inhibitory concentration (MIC) measurements, and bacterial microscopy post-MICs were performed.

**Results:** The assay demonstrated robust performance, evidenced by high Z’-factor and signal-to-noise (S/N ratios. Screening identified 57 active compounds (1.2% of the library), including *β*-lactams and three non-antibiotic molecules—suloctidil, isorotenone, and alexidine—that exhibited concentration-dependent antibacterial activity. Alexidine showed the most potent activity, with the lowest K_50_ (2.7×10^−3^ mM) and MICs of 0.004 mM for *E. coli* and 0.015 mM for *P. aeruginosa*. Suloctidil and isorotenone induced spherical cell morphology, while alexidine induced a filamentous phenotype, indicative of envelope disruption. The assay also identified antibiotics for monotherapy and combination therapy, with ampicillin, alexidine, and suloctidil enhancing chloramphenicol’s efficacy against *E. coli* MG1655.

**Conclusions:** The LacZ/CPRG reporter assay effectively identified compounds targeting bacterial envelopes, including novel molecules with antibacterial activity against GN pathogens, making it a promising tool for antibiotic discovery or combination therapy.

## Introduction

Infectious diseases caused by pathogens remain a major global health threat, with 1.5 billion infections and 4.6 million deaths annually^1–3^. The rise of multidrug-resistant organisms (MDROs), particularly Gram-negative bacteria (GNB), is a critical global challenge^1, 3, 4^. These pathogens’ intrinsic resistance complicates treatment and drives high morbidity and mortality rates ^5^. MDROs caused 700,000 deaths in 2014, with projections reaching 10 million annually by 2050 ^6^. In the U.S., MDROs account for 160,000 deaths each year, while Europe reports 138,000, highlighting the urgent need for effective intervention^7–9^. MDRO-related healthcare costs exceed $35 billion annually in the U.S.^10^, with a global economic burden approaching $100 trillion by mid-century^6^. Healthcare-associated infections (HAIs) add significant economic strain, with U.S. expenditures between $7.2 and $14.9 billion in 2016^11^.

GNB cause approximately half of all bacterial infections, with higher mortality rates compared to Gram-positive (GP) organisms^10^. Treating multidrug-resistant GNB infections, such as sepsis and pneumonia, is challenging due to the outer membrane acting as a barrier to antibiotics and mediating intrinsic resistance^12^. Antimicrobial resistance (AMR) in GNB results from antibiotic-inactivating enzymes or nonenzymatic mechanisms, which can be intrinsic or acquired via mutations or gene transfer^13, 14^. The global spread of MDROs, driven by AMR, has hindered the management of carbapenem-resistant *Enterobacteriaceae*, like *Klebsiella pneumoniae* and *Escherichia coli*^4, 5, 15^.

Therapeutic options and understanding of resistance mechanisms for GNB infections remain limited^16^. Advances in GNB envelope biogenesis studies^16, 17^ highlight the need for innovative antibiotic susceptibility test (AST) methods, narrow-spectrum drugs, and diagnostics^5^. AST technologies include phenotypic and genotypic approaches, such as conventional, automated, and high-throughput screening (HTS), but traditional culture-based AST remains labor-intensive and time-consuming^18^. Recent innovations in this area include Rapid Electrochemical-Based PCR-Less^19^, and photonic-based methods^20^, imaging-based AST^21, 22^, and Raman scattering microscopy^23^. Molecular approaches such as whole-genome sequencing^24^, microfluidics^22, 25^, proteomics^26^, and machine learning-assisted AST^27, 28^ can enhance resistance detection in GN and GP bacteria^28^. Emerging technologies, such as nanoliter droplets, digital imaging, and HTS, target bacterial growth and whole-cell assays^29–34^ can provide new angles for research, although drug discovery now focuses on cellular processes^35, 36^, leveraging cell wall reporter assays^37^ and biochemical screens^35, 38–41^, targeting novel cell wall or membrane inhibition mechanisms^42^.

Despite advancements in diagnostic tools for antibacterial resistance ^28, 43–46^, universal platforms remain elusive due to complexity, cost, and challenges in targeting the GNB outer membrane^24^. With AMR on the rise and 50% of antibiotic treatments initiated without accurate pathogen identification or appropriate antibiotics^47^, there is an urgent need for high-throughput methods targeting GN bacterial envelopes to identify novel, small, and effective molecules.

Here, we present a novel biochemical HTS assay that detects small molecules disrupting GN envelope integrity, diverging from traditional viability-based screens. This LacZ/CPRG-based method, optimized in liquid Luria-Bertani (LB) culture, detects envelope permeabilisation in *E. coli* MG1655, a GN model organism, adapting Paradis-Bleau et al.’s agar-based assay^16^. In the CPRG assay, absorbance at 575 nm measures *β*-galactosidase activity, which serves as a proxy for bacterial viability, membrane integrity, or metabolic activity. Utilising *β*-galactosidase (LacZ) to cleave CPRG, the method produces a measurable colour change upon envelope disruption, enhancing throughput for efficient antibacterial screening. Since the assay indirectly measures CPRG hydrolysis rather than directly assessing bacterial viability or antibiotic uptake, the absorbance signal reflects whether bacteria remain metabolically active despite exposure to the compound or antibiotic. We validated this assay using pathogenic strains, including *E. coli* and *Pseudomonas aeruginosa*, and successfully screened the LOPAC^1280^ and KD2^4761^ libraries^16, 17, 29^. Our assay identified several active compounds, including *β*-lactams, suloctidil, and alexidine, which disrupt bacterial envelope integrity, highlighting their potential for monotherapy and combination therapy.

## Materials and Methods

### Bacterial Strains and Growth Conditions

The *E. coli* strain MG1655 (rfb-50, O-antigen defective)^48^ was employed for HTS optimisation. Assay validation was conducted using Enterohemorrhagic *E. coli* (EHEC, EDL933, O157:H7), Enteropathogenic *E. coli* (EPEC, E2348/69, O127:H6Str)^49^, Uropathogenic *E. coli* (UPEC, CFT073, O6:K2:H1)^50^, and pathogenic *P. aeruginosa* strain PAO1, representing clinically relevant GN pathogens. *E. coli* and *P. aeruginosa* strains were cultured in liquid LB medium (1% tryptone, 0.5% yeast extract, 0.5% NaCl), at 37°C.

### LacZ/CPRG Reporter Assay Optimisation for 96-Well and 384-Well Plates

The LacZ/CPRG phenotypic screen assay, initially developed by Paradis-Bleau et al. for solid media culture to assess cell envelope biogenesis defects^16^, was optimised for automation to screen for small molecules targeting the GN bacterial envelope. Ampicillin (VWR #CA89233-904), a *β*-lactam antibiotic that damages the bacterial envelope^51^, served as the positive control, and chloramphenicol (VWR #CA97061-244) as the negative control^52^. To optimise assay sensitivity, ampicillin, CPRG, and Isopropyl *β*-D-thiogalactopyranoside (IPTG) concentrations were titrated in two separate experiments (Figure S1 and Figure 1a-e). Time courses were evaluated from 3 hours to overnight, with the volume of LB medium reduced to 100 mL for 96-well plates and 55 mL for 384-well plates while still providing robust readout and Z-factor. Lower concentrations of ampicillin and chloramphenicol were used for slower colour development in controls.

**Figure 1.**
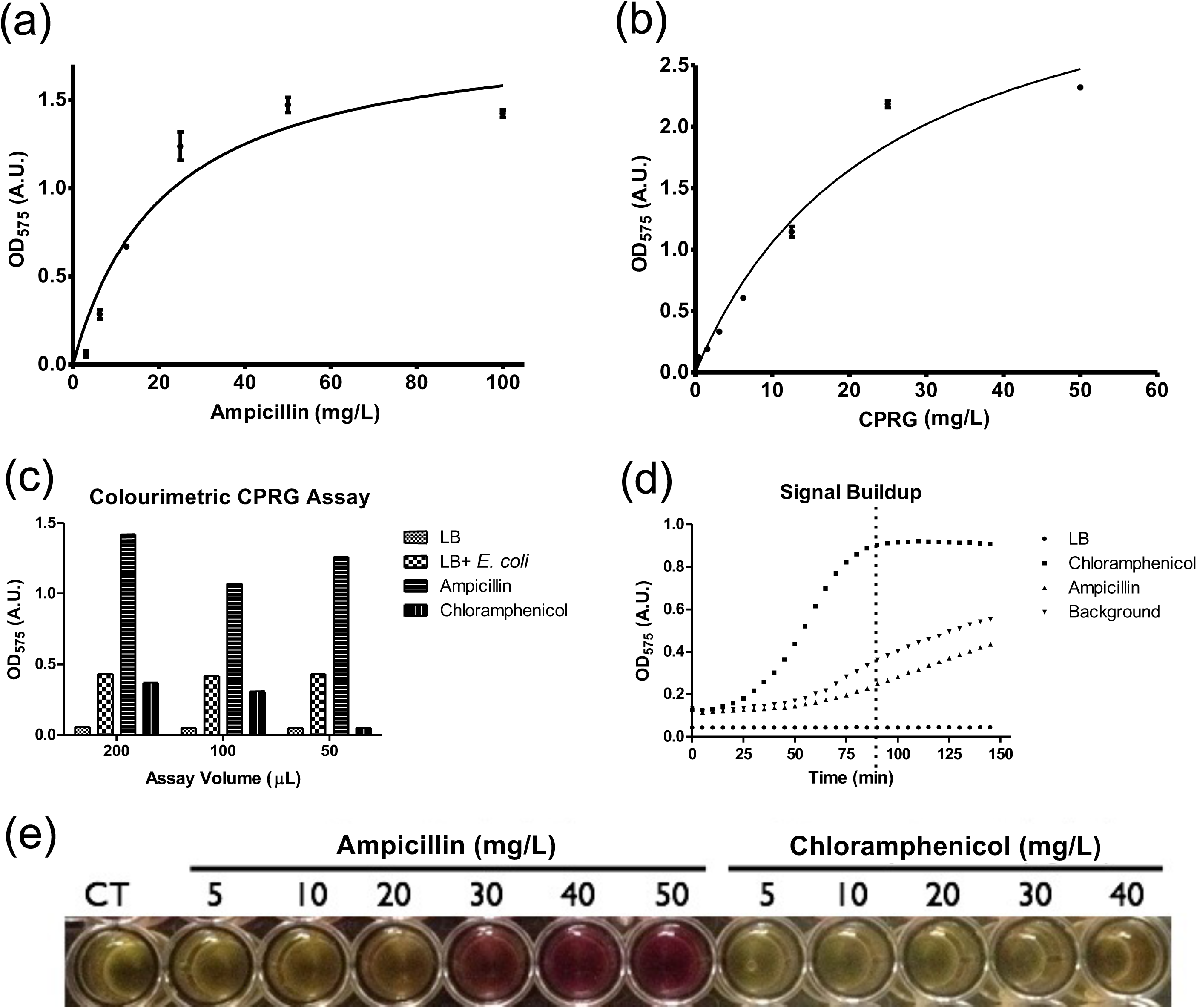
Optimisation of the LacZ/CPRG assay for high-throughput screening in liquid media, measured at OD_575 nm_. Concentration-dependent effects of ampicillin (a) and CPRG (b) were assessed by titration into the reaction mixture in separate experiments in a 50 µL reaction volume. Curves were fitted using the Michaelis-Menten equation, yielding binding constants (Km) of 21 mg/L and 25 mg/L for ampicillin and CPRG, respectively. Colourimetric assay (c) readings after 3 hours of incubation for the negative control (chloramphenicol), positive control (ampicillin), fresh LB medium (to assess potential contamination), and LB with *E. coli* MG1655 culture (to establish background levels). Signal accumulation over time in a 50 µL reaction volume (d), plateauing after approximately 3 hours. Microplate wells demonstrating assay specificity for envelope-targeting antibiotics (e) such as ampicillin, with no response to chloramphenicol, which inhibits protein synthesis. Signals were stable for over 3 hours and exhibited a concentration-dependent response to *β*-lactam ampicillin. Results confirm that concentrations of 40 mg/L ampicillin and 40 mg/L CPRG (chromogenic substrate) generate robust and reproducible signals.

### 96-Well Plate Format Optimisation

For the 96-well plate format (Nunc #167314), a fresh *E. coli* MG1655 colony was inoculated into 4 mL of sterile LB medium supplemented with 100 µM IPTG (VWR #71004-004) (4 µL of the 0.1 M IPTG stock solution) in a test tube and incubated overnight at 37°C with shaking (250 rpm). The overnight culture was diluted 1:50 by adding 500 µL to 25 mL of sterile LB in 250 mL Erlenmeyer glass flask supplemented with 100 µM IPTG (25 µL of the 0.1 M IPTG stock solution) and incubated at 37°C with shaking until mid-log phase (OD_600nm_ = 0.5, approximately 90 minutes). CPRG (VWR #CA80051-016) was added to the culture to a final concentration of 40 mg/L (from the 20 mg/mL stock solution). The assay was performed by transferring 200 µL of CPRG-containing culture into each well of the 96-well plate. Bacterial cells alone were used as the blank, and LB medium was included in some wells to assess potential contamination. In select wells, 40 mg/L ampicillin (positive control) and 40 mg/L chloramphenicol (negative control) were added (4 µL of the 2 mg/mL stock solutions). The plates were incubated at room temperature with shaking. Colour development was assessed at OD_575nm_ using a microplate reader (PowerWave XS, Biotek). Assay quality, consistency, and sensitivity were evaluated by determining Z’ (Z-prime), Z-factor, and signal-to-noise (S/N) ratios^53^.

### 384-Well Plate Format for High-Throughput Screening

For automation and high-throughput screening, the assay was scaled down to 384-well plates (Thermo Fisher #3701), following the same procedure unless otherwise stated. The overnight culture was diluted 1:50 by adding 2 mL of culture to 100 mL of sterile LB supplemented with 100 µM IPTG in 2 L flask and incubated at 37°C with shaking until OD_600nm_ reached 0.6 (approximately 2 hours). Two 384-well plates were prepared, with each well receiving 25 µL of LB medium and specific compounds. In Plate 1, half of the wells were treated with 70 nL of ampicillin (30 mg/mL in DMSO), while the other half received 70 nL of DMSO as a negative control (AMP-). To assess potential contamination, 50 µL of fresh LB was added to all wells in column 24. In Plate 2, half of the wells contained 70 nL of ampicillin (30 mg/mL in DMSO), while the remaining wells contained 70 nL of chloramphenicol (30 mg/mL in DMSO). A 30 mL bacterial culture was prepared and supplemented with CPRG to an initial concentration of 80 mg/L. A Thermo Combi was used to transfer 25 µL of the culture into each well of both plates, ensuring a final CPRG concentration of 40 mg/L (Figure S2). The plates were incubated at room temperature on a rotating platform with continuous agitation. After 3 hours, OD_575nm_ was measured using a microplate reader (PowerWave XS, Biotek). To evaluate whether compounds promoted slow leakage of the cell wall, OD was recorded again after overnight incubation and compared to the values after 3 hours. Z’ values, S/B, and S/N ratios were calculated to assess the quality of the screening assay^53^. The assay was also performed on pathogenic *E. coli* strains (EHEC EDL933, O157:H7; EPEC E2348/69, O127:H6; UPEC CFT073, O6: K2:H1)^49, 50^ and compared with strain MG1655 (rfb-50, O-antigen defective)^48^.

### Automation of the CPRG Assay by Screening the LOPAC^1280^ Library

The LOPAC^1280^ library (4 plates, Sigma), comprising well-characterised compounds with known activities, was utilised to assess the specificity and sensitivity of the automated process. Culture preparation, bacterial inoculation, and incubation steps followed the protocol described in the section on the adaptation of the colourimetric CPRG assay in a 384-well format. Compounds from the LOPAC^1280^ library were pin transferred using a PlateMate Plus and FP3 pin tool, achieving a final concentration of about 6.0 µM/well. For each plate, 30 mL of bacterial culture was supplemented with CPRG (final concentration, 80 mg/L) and ampicillin (final concentration, 40 mg/L) and using a dispenser, 25 µL of bacterial culture containing CPRG and ampicillin was transferred to the wells in row 1 of all four plates as a positive control, with row 24 reserved for the negative control solution. A CPRG-bacterial culture (final CPRG concentration, 80 mg/L) was prepared, and 25 µL was transferred into each well of the rest of the plate, with the resulting final CPRG concentration being 40 mg/L. The plates were incubated at room temperature on a plate shaker After 3 hours and overnight incubation, absorbance was measured at OD_575nm_ using a microplate reader (PowerWave XS, Biotek).

### Automation Parameters of the CPRG Assay in HTS Using the KD2^4761^ Library

The KD2^4761^ compound library (16 plates) comprises various subsets, containing compounds with both known and unknown activities. It consists of compounds from the Prestwick (1120; 4 plates), Microsource (2000; 7 plates), Sigma-LOPAC^1280^ (1280; 4 plates), and the BIOMOL (361;1 plate). The procedures followed were those described in the section on screening the LOPAC^1280^ library, with the following modifications. The percentage of inhibition was determined using the following formula: % Inhibition = (sample data / average background data) × 100% - 100%

### Cherry-Picking for Confirmation of 72 Active Envelope-Targeting Compounds

To confirm the actives identified from the LOPAC and KD_2_ library screens, a cherry-picking assay was performed. The procedures described for the LOPAC library were followed, with the following modifications. Compounds (0.5 μL, with a final concentration of 45 μM in each assay well) were cherry-picked by hand from the primary 96-well compound plates to the KD2 cherry-pick assay plates. Absorbance at 575 nm and 660 nm was monitored for all compounds over 11 hours, with 30-minute intervals for the first 3 hours, followed by 1-hour intervals for the remainder of the experiment. Background absorbance of compounds was subtracted from the assay readings before analysis. The percentage of inhibition was calculated as previously described.

### Concentration-Response Curves

To evaluate the activity of four non-antibiotic envelope-targeting compounds (isorotenone, alexidine dihydrochloride, suloctidil, and thiram) in a concentration-dependent manner, concentration-response curves were generated. The procedures described in the LOPAC screening were followed, with additional modifications. The four non-antibiotic compounds were serially diluted and prepared in DMSO, with final concentrations ranging from 25 μM to 50 nM. Absorbance at 575 nm and 660 nm was monitored for all compounds over 20 hours, with 1-hour intervals. Background absorbance was subtracted from the assay readings before calculating the percentage of inhibition.

To confirm the chemistry, bacterial cell wall permeabilization activities, and the concentration-dependent response of these candidate envelope-targeting antibacterials, six compounds were newly purchased (Sigma-Aldrich, Canada): alexidine, suloctidil, isorotenone, and three control compounds (vancomycin, cefaclor, and cefsoludin). Concentration-response curves were generated for these compounds. The procedures and analysis were as described above, with additional modifications.

The six repurchased compounds were prepared as 20 or 25 mM stocks in DMSO. Serial dilutions of these six compounds were prepared in DMSO, with final concentrations ranging from 2.5 mM to 2.5 μM.

### Co-Permeabilisation Assay

To test the activity of chloramphenicol, which does not penetrate the *E. coli* membrane, and its mediation by the cell wall permeabilizing compounds identified in the screen, a co-permeabilization experiment was conducted. The overnight culture was diluted 1:50 by adding 2 mL of culture to 100 mL of sterile LB, supplemented with 100 µM IPTG (100 µL of a 0.1 M IPTG stock solution), and transferred to a 250 mL flask. The flasks were incubated at 37°C with 250 rpm shaking until the OD_600_ nm reached 0.8, typically taking about 2 hours (recording OD began after 30 minutes of incubation). CPRG was supplemented to a final concentration of 40 mg/L, from a stock solution of 20,000 mg/L (20,000 µg/mL). For the assay, 25 µL of CPRG-containing culture and 25 µL of fresh LB were added to each well. Compounds were added from the dilution plate, followed by absorbance readings at OD_575_ nm and OD_660_ nm for 5 hours, with measurements every 45 minutes using the Power Wave plate reader.

### Minimal Inhibitory Concentration (MIC) and Microscopic Assays

MIC and phenotypic assays were conducted in triplicate to evaluate the inhibitory effects of selected compounds—isorotenone, suloctidil, and alexidine—on *E. coli* (strain K12 MG1655) and a pathogenic strain of *P. aeruginosa* PAO1, representing a Gram-negative bacterium. MIC assays were performed in Luria-Bertani (LB) broth containing 1% NaCl and varying concentrations of the test compounds, as indicated. Cultures were initiated with bacterial suspensions at 2×10^6^ CFU/mL and incubated at 37°C for 18 hours. Growth was monitored by measuring optical density at 575 nm at 30-minute intervals. For phenotypic and microscopic analyses, single colonies of *E. coli* and *P. aeruginosa* were cultured overnight in 6 mL of LB medium at 37°C with arbitrary agitation in test tubes. Cultures were diluted to an initial optical density at 600 nm (OD_600_) of 0.02 in fresh LB medium and transferred to 250 mL Erlenmeyer flasks containing 25 mL LB. Incubation was carried out at 37°C with constant agitation at 250 rpm. Cell growth was recorded by periodic OD_600_ measurements. At the mid-log phase (OD_600_ ≈ 0.5), 3 µL of the cultures were spotted onto 0.8% agarose pads for visualisation. Phase-contrast images of the cells were captured using a Nikon Eclipse E600 microscope equipped with a 100× oil immersion objective and a DS-Ri2 Nikon camera.

## Results

### Assay Optimisation for 96- and 384-Well Plates

The CPRG assay for B-gal leakage from bacterial cells was successfully optimised for 96- and 384-well plate formats. Concentration-dependent curves for ampicillin and CPRG were fitted using the Michaelis-Menten equation to evaluate data quality. Binding constants for ampicillin and CPRG were determined to be 21 mg/L and 25 mg/L, respectively, within a 50 µL reaction volume (Figure 1a, b). These results demonstrate that concentrations of 40 mg/L ampicillin (positive control) and 40 mg/L CPRG (chromogenic substrate) yield robust signal intensities. As illustrated in Figure 1c, the assay is readily scalable from 200 µL to 50 µL (from 96 well to 384 well plate) without compromising performance. A fourfold increase in absorbance units (A.U.) was observed in the positive control (ampicillin) relative to the negative control (chloramphenicol) and a threefold increase relative to the background (LB + *E. coli*) (Figure 1c). The absorbance signal, monitored over time, plateaued after approximately 90 minutes of incubation (Figure 1d). Based on these findings, the reaction mixture requires a minimum incubation period of 2 hours before measurement, with 3 hours providing optimal conditions. The assay exhibited excellent performance in liquid media, demonstrated specificity for envelope-targeting antibiotics such as ampicillin, and was well-suited for high-throughput screening applications (Figure 1e).

### Strain Sensitivity and Assay Performance

We aimed to identify small molecules targeting the Gram-negative bacterial envelope by screening libraries using the CPRG assay. This phenotypic assay detects defects in the envelope through restricted LacZ access to extracellular CPRG. A positive result occurs when cell lysis releases LacZ or increased envelope permeability allows CPRG entry^16^ (Figure 2a). The assay, developed using the O-antigen-defective *E. coli* MG1655 strain, was designed to assess whether envelope permeability defects could be transferred to pathogenic *E. coli* strains possessing complete LPS structures. A concentration of 0.01 mM ampicillin was sufficient to generate a positive readout in the assay across all tested strains. Among them, EHEC (enterohaemorrhagic *E. coli*) exhibited the highest sensitivity, albeit with a higher background signal, whereas UPEC (uropathogenic *E. coli*) demonstrated a weaker response (Figure 2b).

**Figure 2.**
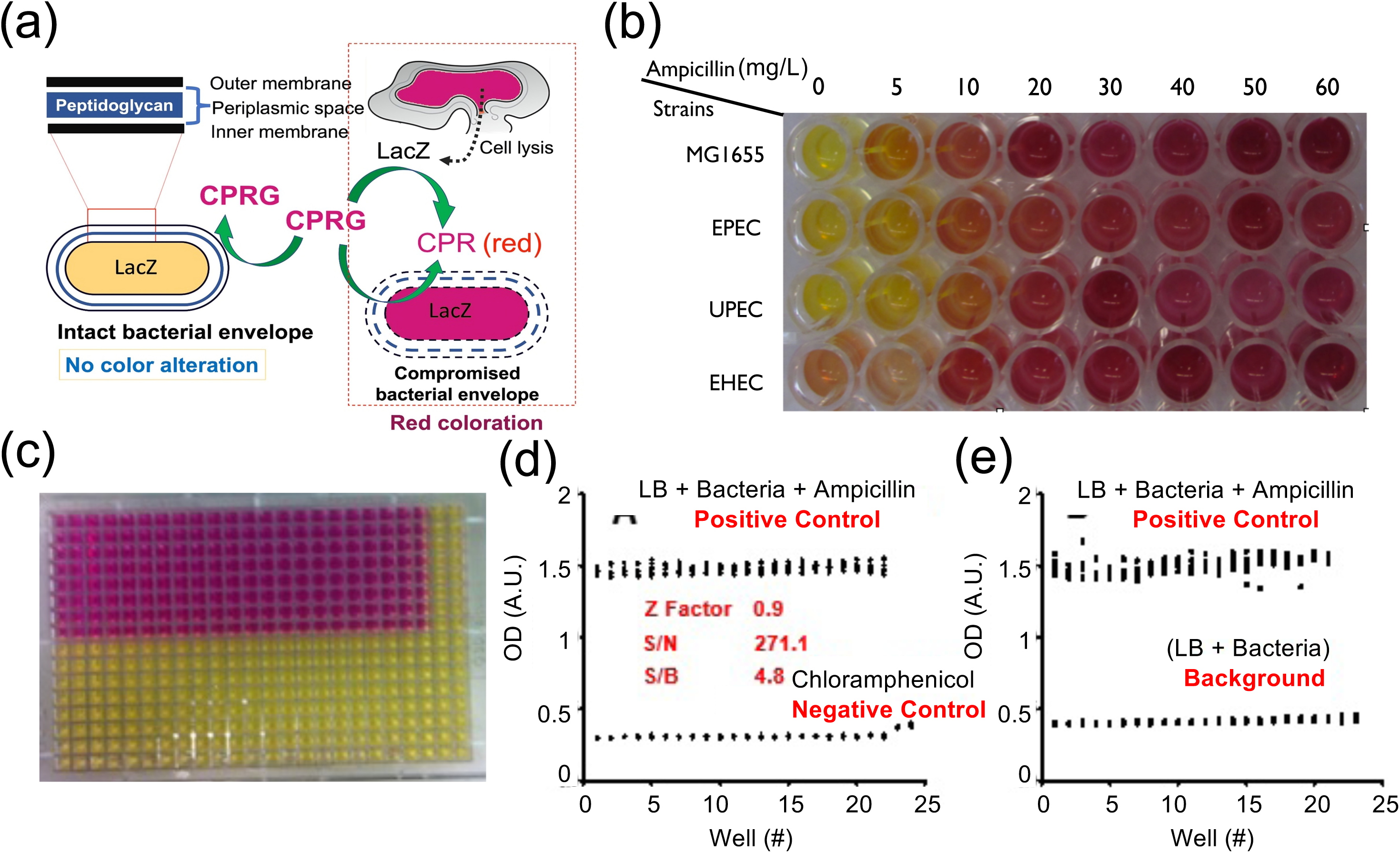
Statistical validation of ampicillin’s impact on envelope permeability and LacZ/CPRG activity in *E. coli* with statistical validation. (a) Fundamental of LacZ/CPRG assay (b) LacZ/CPRG assay demonstrating colour change and envelope permeability effects induced by varying concentrations of ampicillin on pathogenic *E. coli* strains. Spectrophotometric readings were taken at 575 nm. Enterohaemorrhagic (EHEC EDL933, O157:H7), Enteropathogenic (EPEC E2348/69, O127:H6), and Uropathogenic (UPEC CFT073, O6: K2:H1) strains were compared with the O-antigen-defective strain MG1655 (rfb-50). Cells were cultured in LB medium with 100 µM IPTG for LacZ induction at 37°C until OD_600nm_ of 0.5, then 200 µL was transferred to a new and CPRG (40 µg/mL) and ampicillin were added to assess envelope perturbations. After incubation for 30 minutes at room temperature, colour change was recorded. (c) Validation of the LacZ/CPRG HTS assay in a 384-well plate format (55 µL liquid media per well). (d) Statistical analysis and signal separation from ampicillin (positive control) and chloramphenicol (negative control). (e) Signal separation from background and observed colour change for ampicillin. Z’-factor, signal-to-noise (S/N), and signal-to-background (S/B) ratios were used to assess assay robustness. Z’-factor values was 0.9 in 384-well formats.

### Statistical Validation and Assay Robustness

To validate assay performance, we assessed key statistical parameters, including the Z’-factor, Signal-to-Background (S/B) ratio, and Signal-to-Noise (S/N) ratio. These metrics are crucial for determining the assay’s robustness and suitability for high-throughput screening (HTS). We measured the Z’-factor and background signal to evaluate whether compound concentration would promote gradual cell wall leakage over a 3-hour time course and following overnight incubation. The background increased overnight at 40 mg/L CPRG; however, the Z’-factor (0.6) confirmed the assay’s reliability for HTS applications. Based on existing results^16^ and our further analysis, reducing CPRG concentration by 20 mg/L may help mitigate the background signal after 3 hours, if necessary (Table 1). Despite this adjustment, a reduction in CPRG concentration does not enhance overall HTS parameters.

**Table 1.** Statistical factors for 20 mg/L and 40 mg/L CPRG across 3-hour and overnight incubation.

| Incubation | CPRG 20 mg/L | CPRG, 40 mg/L |
| --- | --- | --- |
| 3 hours | Z'- factor = 0.8 | Z'- factor = 0.8 |
|  | S/B = 2.3 | S/B = 3.4 |
|  | S/N = 81 | S/N = 194 |
| Overnight | Z'- factor = 0.1 | Z'- factor = 0.6 |
|  | S/B = 1.4 | S/B = 1.8 |
|  | S/N = 5.9 | S/N = 9.4 |

The assay was initially optimised in a 96-well plate format (100 µL), yielding a Z’-factor of 0.6, an S/B ratio of 3.9, and an S/N ratio of 38.5. Miniaturisation to a 384-well format (55 µL) improved these values to 0.9, 4.8, and 271, respectively (Figure 2c). As the Z’-factor is a key indicator of assay quality, its consistency and sensitivity were further assessed. The Z’-factor quantifies the separation between positive (ampicillin) and negative (chloramphenicol) controls (Figure 2d), while assessing the positive control against the background (LB media + bacteria) also provides insight into the likelihood of false positives or negatives (Figure 2e). According to established guidelines^53^, a Z’-factor above 0.5 indicates minimal sensitivity impact, while values between 0.5 and 1.0 signify an excellent assay.

### Automation by Screening the LOPAC^1280^ Library

The LOPAC^1280^ library, composed of compounds with known biological activities, was used to evaluate automation specificity and sensitivity at room temperature after 3 hours and overnight incubation. The automation test yielded robust data, with an excellent average Z’-factor of 0.6 a strong signal-to-noise (S/N) ratio of 17, and a signal-to-background (S/B) ratio of 2.6. Screening identified five active compounds after 3 hours (Figure 3a) and eleven after overnight incubation (Figure 3b). Data are presented as % inhibition, normalised to controls. A 25% inhibition cut-off was applied, determined by three times the standard deviation above the assay plate mean (Table 2). Among these, ten overlapped with the KD2 screen (Table 3), while trimethoprim, exclusive to the LOPAC library, was uniquely identified.

**Figure 3.**
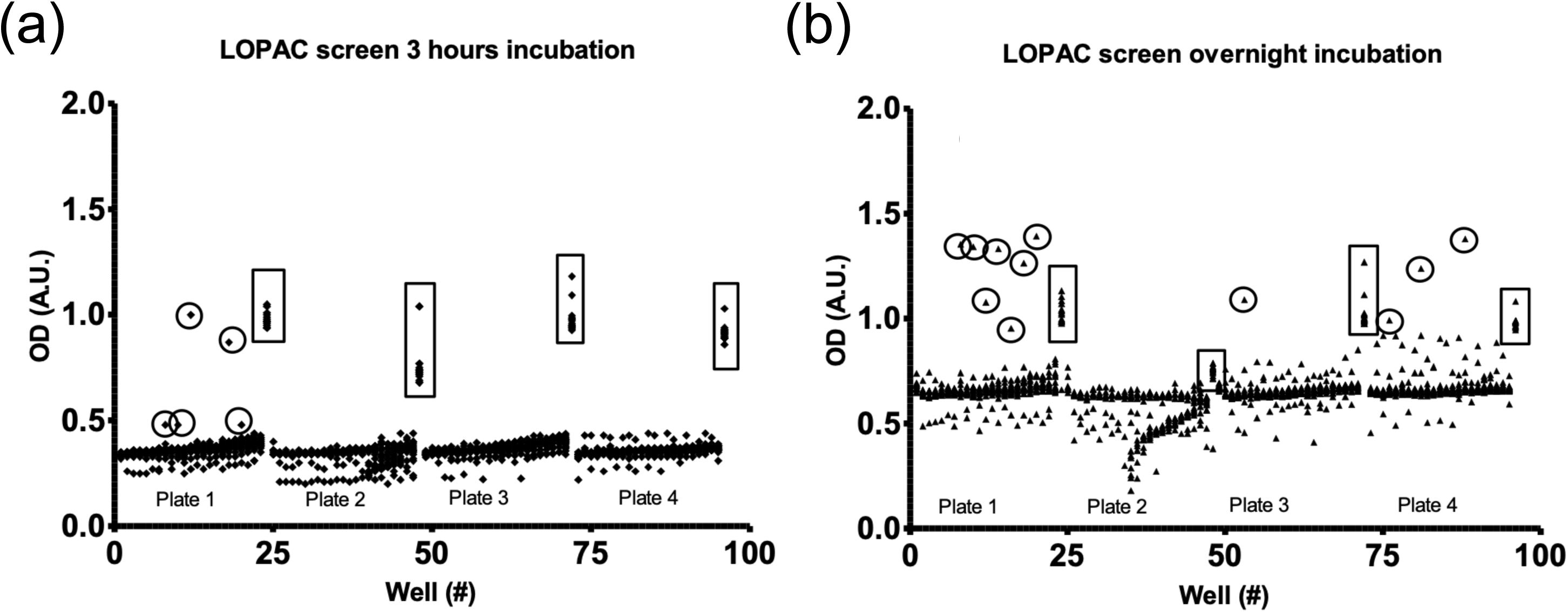
Distribution plot of LOPAC^1280^ compound screening after 3-hour (a) and overnight (b) incubations, measured spectrophotometrically at OD_575_ nm. Positive controls are indicated by boxes, and potential hits by circles. Following normalisation to controls, five compounds from the 3 hr incubation and eleven compounds from the overnight incubation exhibiting >25% inhibition were identified as actives after 3 hours and overnight incubation. The 25% threshold, set at three times the standard deviation above the mean, ensures a robust cutoff for identifying significant effects (see Table 2). The chart displays data points where the X-axis represents well numbers (#) across 384-well plates (16 rows × 24 columns), and the Y-axis shows optical density (OD) values.

**Table 2.** Active compounds from the LOPAC^1280^ screen with >25% OD absorbance.

| Compound name | Function | Group |
| --- | --- | --- |
| Ceftriaxone sodium | Cell wall synthesis | Cephalosporin, 3<sup>rd</sup> gen |
| Cefmetazole sodium | Cell wall synthesis | Cephalosporin, 2<sup>nd</sup> gen |
| Cefazolin sodium | Cell wall synthesis | Cephalosporin, 1<sup>st</sup> gen |
| Cefotaxime sodium | Cell wall synthesis | Cephalosporin, 3<sup>rd</sup> gen |
| Cephapirin sodium | Cell wall synthesis | Cephalosporin, 1<sup>st</sup> gen |
| Cephalothin sodium | Cell wall synthesis | Cephalosporin, 1<sup>st</sup> gen |
| Cefaclor | Cell wall synthesis | Cephalosporin, 2<sup>nd</sup> gen |
| Vancomycin | Cell wall synthesis | Glycopeptides antibiotics |
| Minocycline hydrochloride | Protein synthesis, Cell cycle | Tetracycline/Polyketide |
| Tetraethyl thiuram disulfide | Alcohol dehydrogenase inhibitor | Organosulfur compounds |
| Trimethoprim | Dihydrofolate reductase | Diaminopyrimidine |

**Table 3.**
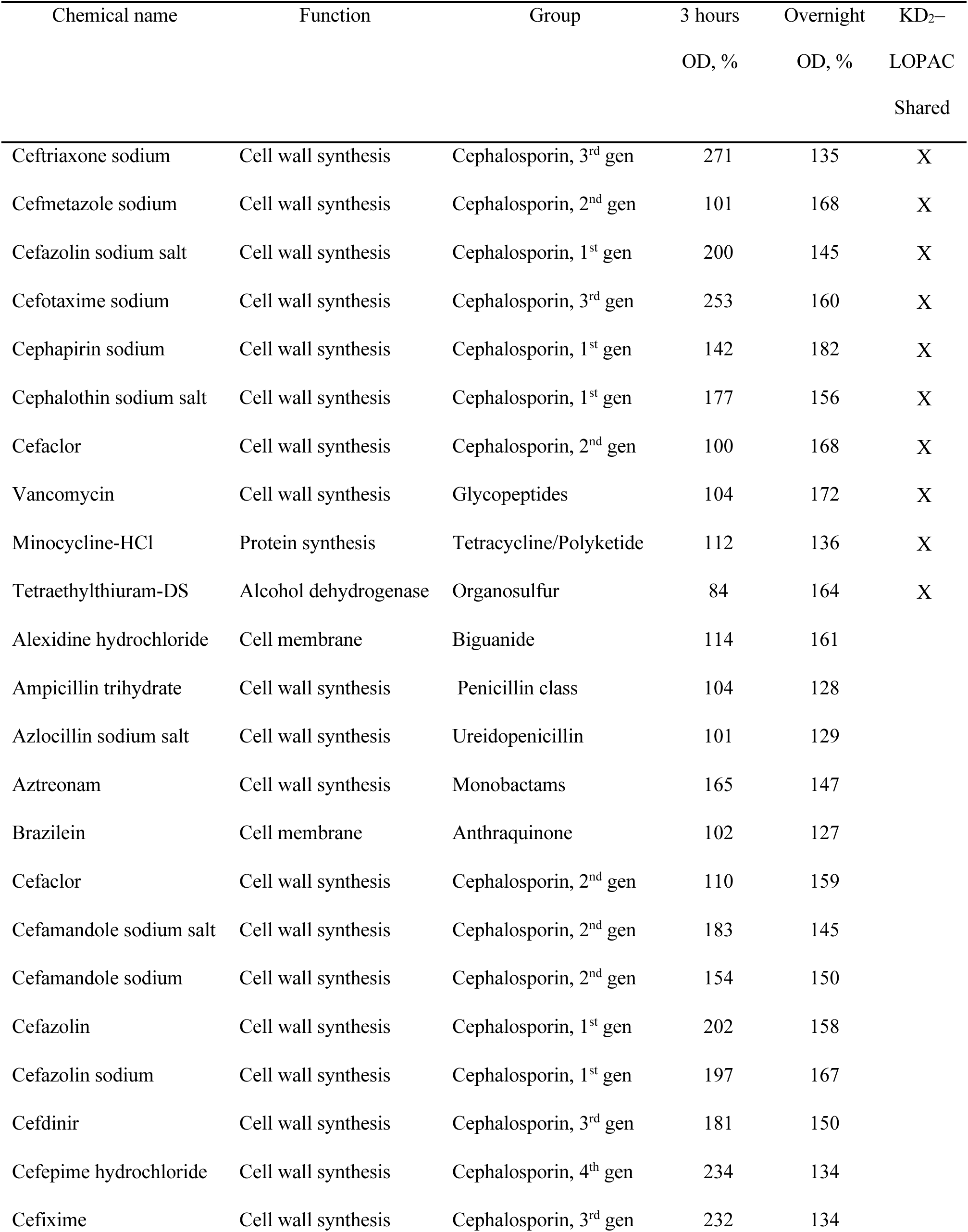
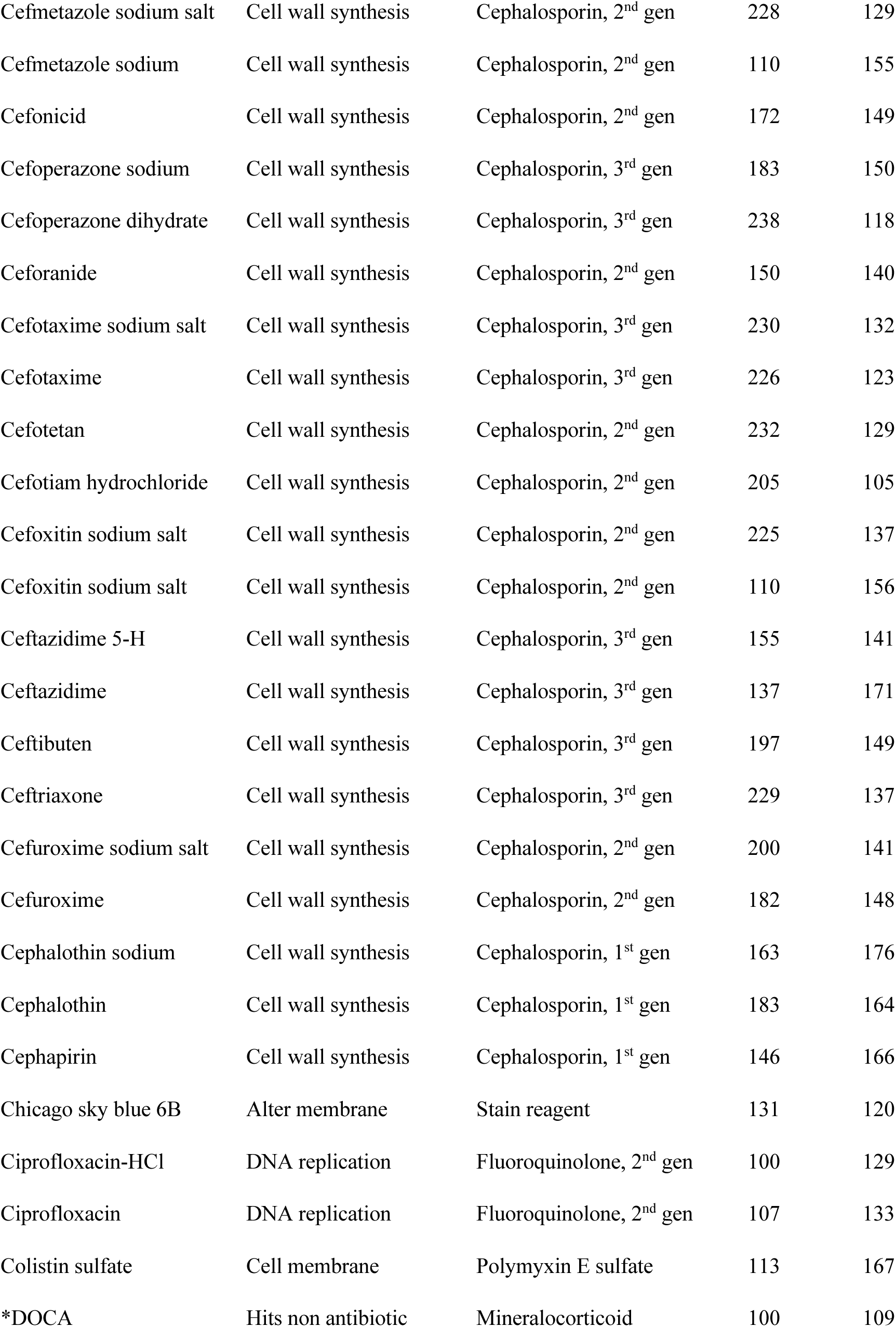
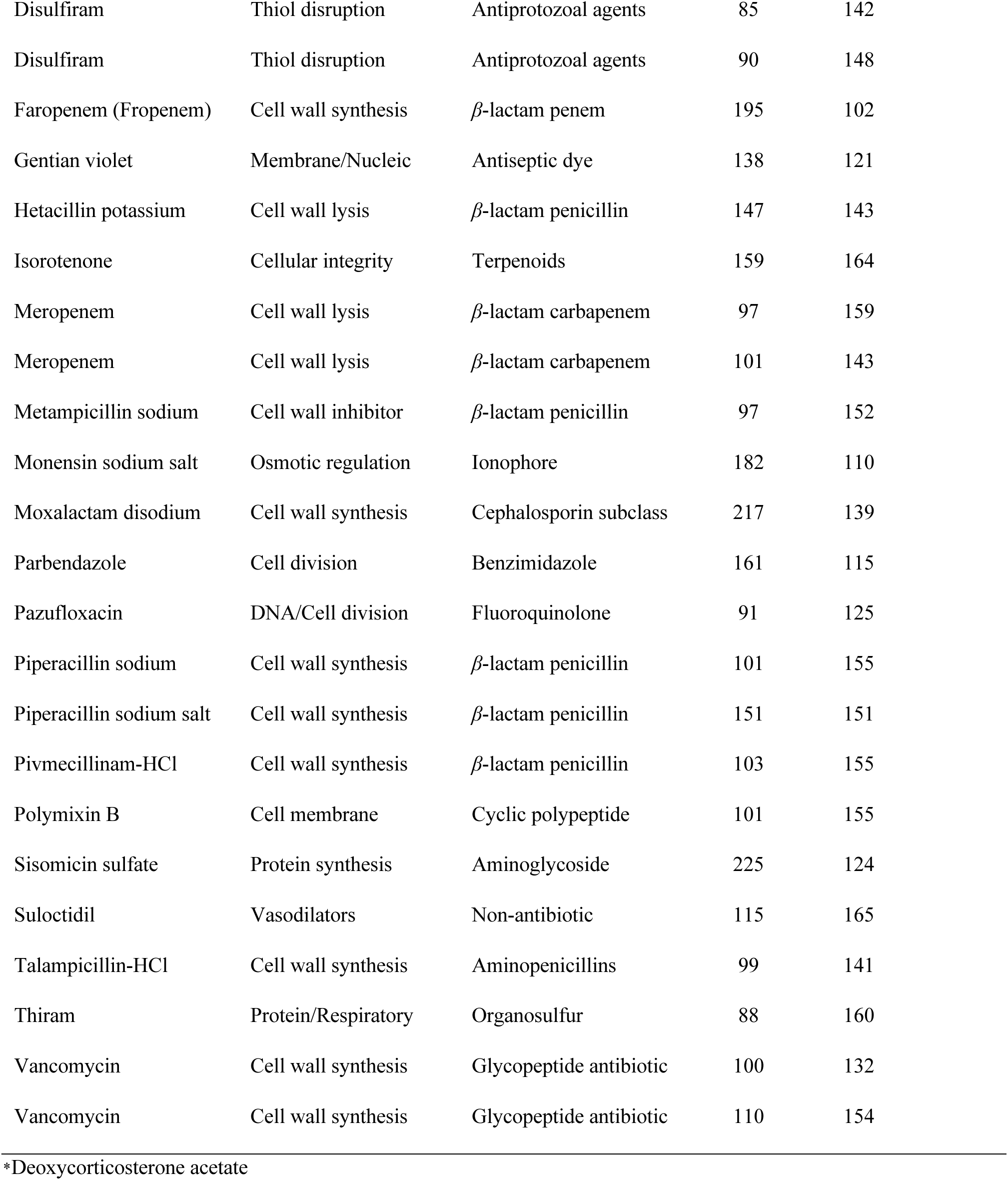
Actives identified from KD_2_^3840^ library screen after normalisation. HTS identified 72 actives (>25% inhibition), with 42 hits at 3 hours and 30 from overnight incubation. Of 11 LOPAC^1280^ hits, 10 were redundant in KD_2_^3840^.

| Chemical name | Function | Group | 3 hours<br>OD, % | Overnight<br>OD, % | KD<sub>2</sub>–<br>LOPAC<br>Shared |
| --- | --- | --- | --- | --- | --- |
| Ceftriaxone sodium | Cell wall synthesis | Cephalosporin, 3<sup>rd</sup> gen | 271 | 135 | X |
| Cefmetazole sodium | Cell wall synthesis | Cephalosporin, 2<sup>nd</sup> gen | 101 | 168 | X |
| Cefazolin sodium salt | Cell wall synthesis | Cephalosporin, 1<sup>st</sup> gen | 200 | 145 | X |
| Cefotaxime sodium | Cell wall synthesis | Cephalosporin, 3<sup>rd</sup> gen | 253 | 160 | X |
| Cephapirin sodium | Cell wall synthesis | Cephalosporin, 1<sup>st</sup> gen | 142 | 182 | X |
| Cephalothin sodium salt | Cell wall synthesis | Cephalosporin, 1<sup>st</sup> gen | 177 | 156 | X |
| Cefaclor | Cell wall synthesis | Cephalosporin, 2<sup>nd</sup> gen | 100 | 168 | X |
| Vancomycin | Cell wall synthesis | Glycopeptides | 104 | 172 | X |
| Minocycline-HCl | Protein synthesis | Tetracycline/Polyketide | 112 | 136 | X |
| Tetraethylthiuram-DS | Alcohol dehydrogenase | Organosulfur | 84 | 164 | X |
| Alexidine hydrochloride | Cell membrane | Biguanide | 114 | 161 |  |
| Ampicillin trihydrate | Cell wall synthesis | Penicillin class | 104 | 128 |  |
| Azlocillin sodium salt | Cell wall synthesis | Ureidopenicillin | 101 | 129 |  |
| Aztreonam | Cell wall synthesis | Monobactams | 165 | 147 |  |
| Brazilein | Cell membrane | Anthraquinone | 102 | 127 |  |
| Cefaclor | Cell wall synthesis | Cephalosporin, 2<sup>nd</sup> gen | 110 | 159 |  |
| Cefamandole sodium salt | Cell wall synthesis | Cephalosporin, 2<sup>nd</sup> gen | 183 | 145 |  |
| Cefamandole sodium | Cell wall synthesis | Cephalosporin, 2<sup>nd</sup> gen | 154 | 150 |  |
| Cefazolin | Cell wall synthesis | Cephalosporin, 1<sup>st</sup> gen | 202 | 158 |  |
| Cefazolin sodium | Cell wall synthesis | Cephalosporin, 1<sup>st</sup> gen | 197 | 167 |  |
| Cefdinir | Cell wall synthesis | Cephalosporin, 3<sup>rd</sup> gen | 181 | 150 |  |
| Cefepime hydrochloride | Cell wall synthesis | Cephalosporin, 4<sup>th</sup> gen | 234 | 134 |  |
| Cefixime | Cell wall synthesis | Cephalosporin, 3<sup>rd</sup> gen | 232 | 134 |  |

|  |  |  |  |  |
| --- | --- | --- | --- | --- |
| Cefmetazole sodium salt | Cell wall synthesis | Cephalosporin, 2<sup>nd</sup> gen | 228 | 129 |
| Cefmetazole sodium | Cell wall synthesis | Cephalosporin, 2<sup>nd</sup> gen | 110 | 155 |
| Cefonicid | Cell wall synthesis | Cephalosporin, 2<sup>nd</sup> gen | 172 | 149 |
| Cefoperazone sodium | Cell wall synthesis | Cephalosporin, 3<sup>rd</sup> gen | 183 | 150 |
| Cefoperazone dihydrate | Cell wall synthesis | Cephalosporin, 3<sup>rd</sup> gen | 238 | 118 |
| Ceforanide | Cell wall synthesis | Cephalosporin, 2<sup>nd</sup> gen | 150 | 140 |
| Cefotaxime sodium salt | Cell wall synthesis | Cephalosporin, 3<sup>rd</sup> gen | 230 | 132 |
| Cefotaxime | Cell wall synthesis | Cephalosporin, 3<sup>rd</sup> gen | 226 | 123 |
| Cefotetan | Cell wall synthesis | Cephalosporin, 2<sup>nd</sup> gen | 232 | 129 |
| Cefotiam hydrochloride | Cell wall synthesis | Cephalosporin, 2<sup>nd</sup> gen | 205 | 105 |
| Cefoxitin sodium salt | Cell wall synthesis | Cephalosporin, 2<sup>nd</sup> gen | 225 | 137 |
| Cefoxitin sodium salt | Cell wall synthesis | Cephalosporin, 2<sup>nd</sup> gen | 110 | 156 |
| Ceftazidime 5-H | Cell wall synthesis | Cephalosporin, 3<sup>rd</sup> gen | 155 | 141 |
| Ceftazidime | Cell wall synthesis | Cephalosporin, 3<sup>rd</sup> gen | 137 | 171 |
| Ceftibuten | Cell wall synthesis | Cephalosporin, 3<sup>rd</sup> gen | 197 | 149 |
| Ceftriaxone | Cell wall synthesis | Cephalosporin, 3<sup>rd</sup> gen | 229 | 137 |
| Cefuroxime sodium salt | Cell wall synthesis | Cephalosporin, 2<sup>nd</sup> gen | 200 | 141 |
| Cefuroxime | Cell wall synthesis | Cephalosporin, 2<sup>nd</sup> gen | 182 | 148 |
| Cephalothin sodium | Cell wall synthesis | Cephalosporin, 1<sup>st</sup> gen | 163 | 176 |
| Cephalothin | Cell wall synthesis | Cephalosporin, 1<sup>st</sup> gen | 183 | 164 |
| Cephapirin | Cell wall synthesis | Cephalosporin, 1<sup>st</sup> gen | 146 | 166 |
| Chicago sky blue 6B | Alter membrane | Stain reagent | 131 | 120 |
| Ciprofloxacin-HCl | DNA replication | Fluoroquinolone, 2<sup>nd</sup> gen | 100 | 129 |
| Ciprofloxacin | DNA replication | Fluoroquinolone, 2<sup>nd</sup> gen | 107 | 133 |
| Colistin sulfate | Cell membrane | Polymyxin E sulfate | 113 | 167 |
| *DOCA | Hits non antibiotic | Mineralocorticoid | 100 | 109 |
| Disulfiram | Thiol disruption | Antiprotozoal agents | 85 | 142 |
| Disulfiram | Thiol disruption | Antiprotozoal agents | 90 | 148 |
| Faropenem (Fropenem) | Cell wall synthesis | β-lactam penem | 195 | 102 |
| Gentian violet | Membrane/Nucleic | Antiseptic dye | 138 | 121 |
| Hetacillin potassium | Cell wall lysis | β-lactam penicillin | 147 | 143 |
| Isorotenone | Cellular integrity | Terpenoids | 159 | 164 |
| Meropenem | Cell wall lysis | β-lactam carbapenem | 97 | 159 |
| Meropenem | Cell wall lysis | β-lactam carbapenem | 101 | 143 |
| Metampicillin sodium | Cell wall inhibitor | β-lactam penicillin | 97 | 152 |
| Monensin sodium salt | Osmotic regulation | Ionophore | 182 | 110 |
| Moxalactam disodium | Cell wall synthesis | Cephalosporin subclass | 217 | 139 |
| Parbendazole | Cell division | Benzimidazole | 161 | 115 |
| Pazufloxacin | DNA/Cell division | Fluoroquinolone | 91 | 125 |
| Piperacillin sodium | Cell wall synthesis | β-lactam penicillin | 101 | 155 |
| Piperacillin sodium salt | Cell wall synthesis | β-lactam penicillin | 151 | 151 |
| Pivmecillinam-HCl | Cell wall synthesis | β-lactam penicillin | 103 | 155 |
| Polymixin B | Cell membrane | Cyclic polypeptide | 101 | 155 |
| Sisomicin sulfate | Protein synthesis | Aminoglycoside | 225 | 124 |
| Suloctidil | Vasodilators | Non-antibiotic | 115 | 165 |
| Talampicillin-HCl | Cell wall synthesis | Aminopenicillins | 99 | 141 |
| Thiram | Protein/Respiratory | Organosulfur | 88 | 160 |
| Vancomycin | Cell wall synthesis | Glycopeptide antibiotic | 100 | 132 |
| Vancomycin | Cell wall synthesis | Glycopeptide antibiotic | 110 | 154 |

\*Deoxycorticosterone acetate

Eight compounds affected cell wall synthesis, while trimethoprim, disulfiram, and minocycline hydrochloride did not but may exert intermediate effects on bacterial cells. Trimethoprim inhibits dihydrofolate reductase, indirectly impairing DNA synthesis^54^. Though it does not interact directly with CPRG, its impact on growth, enzyme expression, or resistance may influence assay results. Disulfiram, used to treat alcohol use disorder, also shows therapeutic potential against Lyme disease, cancer, bacterial infections, and parasitic conditions^55^. Minocycline hydrochloride, a semi-synthetic tetracycline, inhibits protein synthesis in both GP and GN bacteria and exhibits additional non-antibiotic activities^56^. Although these compounds do not directly disrupt cell wall synthesis, their effects on bacterial metabolism, DNA replication, and protein synthesis may indirectly influence CPRG assay outcomes by modulating *β*-galactosidase activity and CPRG hydrolysis.

### Automation by Screening the KD2^4761^ Library in HTS

To evaluate antibacterial activity, we screened 3,840 compounds from the KD2^4761^ library and analyzed their distribution (Figure 4a_1_). The assay identified 72 envelope-targeting candidates exhibiting over 25% inhibition—42 actives at 3 hours and 30 after overnight incubation, indicated by the dotted line (Figure 4a_2_). The screen demonstrated strong assay performance, yielding satisfactory Z’, S/B, and S/N values^53^ (Figure 4b–d). Several active compounds were also present in the KD2^3840^ library (Table 3). Notably, 90% of the identified compounds were known antibiotics, with *β*-lactams targeting cell wall synthesis comprising 80% (57 hits), of which 60% were unique and the remainder duplicates.

**Figure 4.**
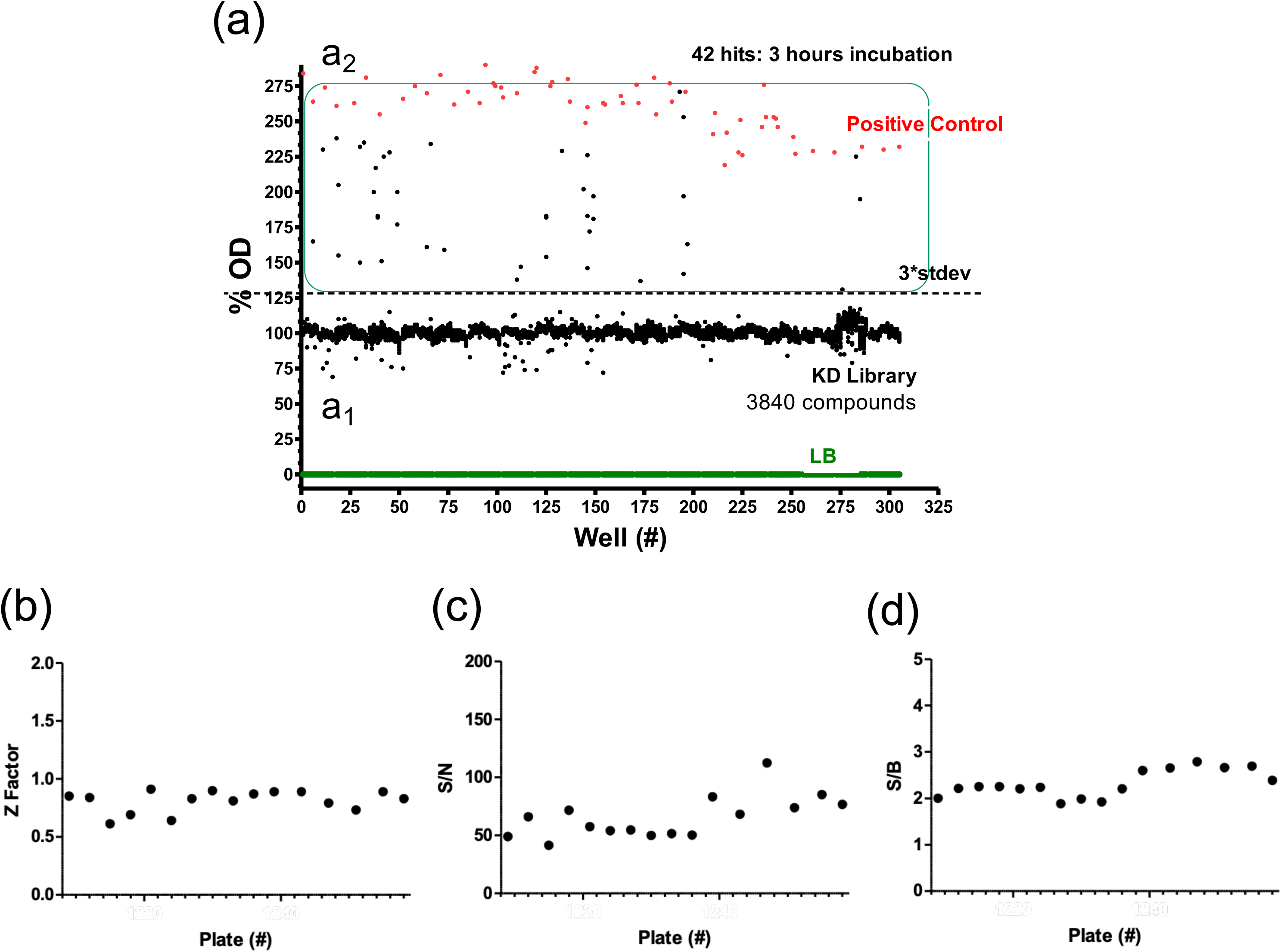
The library screen of 3,840 compounds from KD2-4761 (4a_1_), identifying 72 hits with OD_575_ ≥125% (42 after 3 hours, 30 after overnight incubation). A subset of 42 actives with >25% inhibition after 3 hours is marked by dots (4a_2_), with the dotted line indicating the 25% inhibition threshold, suggesting envelope permeabilisation exceeding 25% relative to the control. Data were normalised to LB alone (0% OD) and a negative control (100% OD). The positive control (ampicillin) had an average OD_575_ of 240% (red dot), with a screen standard deviation of 101 ± 5.7. The LOPAC^1280^ screen identified 11 actives, 10 of which were confirmed in the KD2 screen (Table 2). The chart displays data points where the X-axis represents well numbers (#) across 384-well plates (16 rows × 24 columns), and the Y-axis shows optical density (OD) values. The distribution of Z-Factors (b), S/N (c), and S/B (d) across 16 KD2 plates is shown, with each data point representing a plate on the X-axis.

### Confirmation of 72 Envelope-Targeting Actives by Absorbance Assays

After screening the LOPAC and KD2 libraries using the adapted and optimized CPRG assay for potential envelope-targeting agents, we selected 72 compounds for re-evaluation. Cherry-picking was performed following the protocol outlined in Materials and Methods, with assessments at 3-hour and overnight incubations. Absorbance at 575 nm was monitored over 11 hours to distinguish compounds inducing slow leakage from those exhibiting rapid activity (Figure S3). Each compound produced a distinct “leakage” curve. Thirteen compounds were classified as “slow leakers,” nine as “moderate rate leakers,” and the remainder exhibited rapid action. Seven actives failed to confirm, and eight compounds inhibited bacterial growth without generating a significant increase in absorbance at 575 nm compared to the background and negative controls. Of the 72 compounds screened from KD2^3840^, 57 (∼80%) displayed confirmed *β*-lactams activity, increasing absorbance at 575 nm by >25% at 0.0063 mM (6.3 µM), thereby validating their permeabilising effect. Active compounds comprised 1.2% of the library. Among these, four—isorotenone, alexidine dihydrochloride, suloctidil, and thiram—were non-antibiotic compounds.

### Concentration-Response and Evaluation of Permeabilisation

Interference and artefacts from fluorescence and light detection are common in HTS assays. To address this, primary-screening actives were retested in a concentration-dependent assay to confirm or reject putative actives^57^. We then assessed the envelope-targeting activity of six selected compounds through concentration-response analysis, determining the relationship between compound concentration and permeabilisation. The Permeabilisation constant (K_50_) was calculated as the concentration at which 50% of the maximum permeabilisation effect occurred. Concentration-response assays were performed at room temperature, and curves generated for three antibiotic controls (vancomycin, cefaclor, and cefsoludin), followed by three non-antibiotic compounds (isorotenone, alexidine dihydrochloride, and suloctidil) (Figure 5), with DMSO included as a control, as described in Materials and Methods. Thiram, a strong hit, was excluded from further testing and not analysed in a concentration-dependent manner. This compound, commonly used as a fungicide and ectoparasiticide to prevent fungal diseases in seeds and crops, was not pursued due to its specific application.

**Figure 5.**
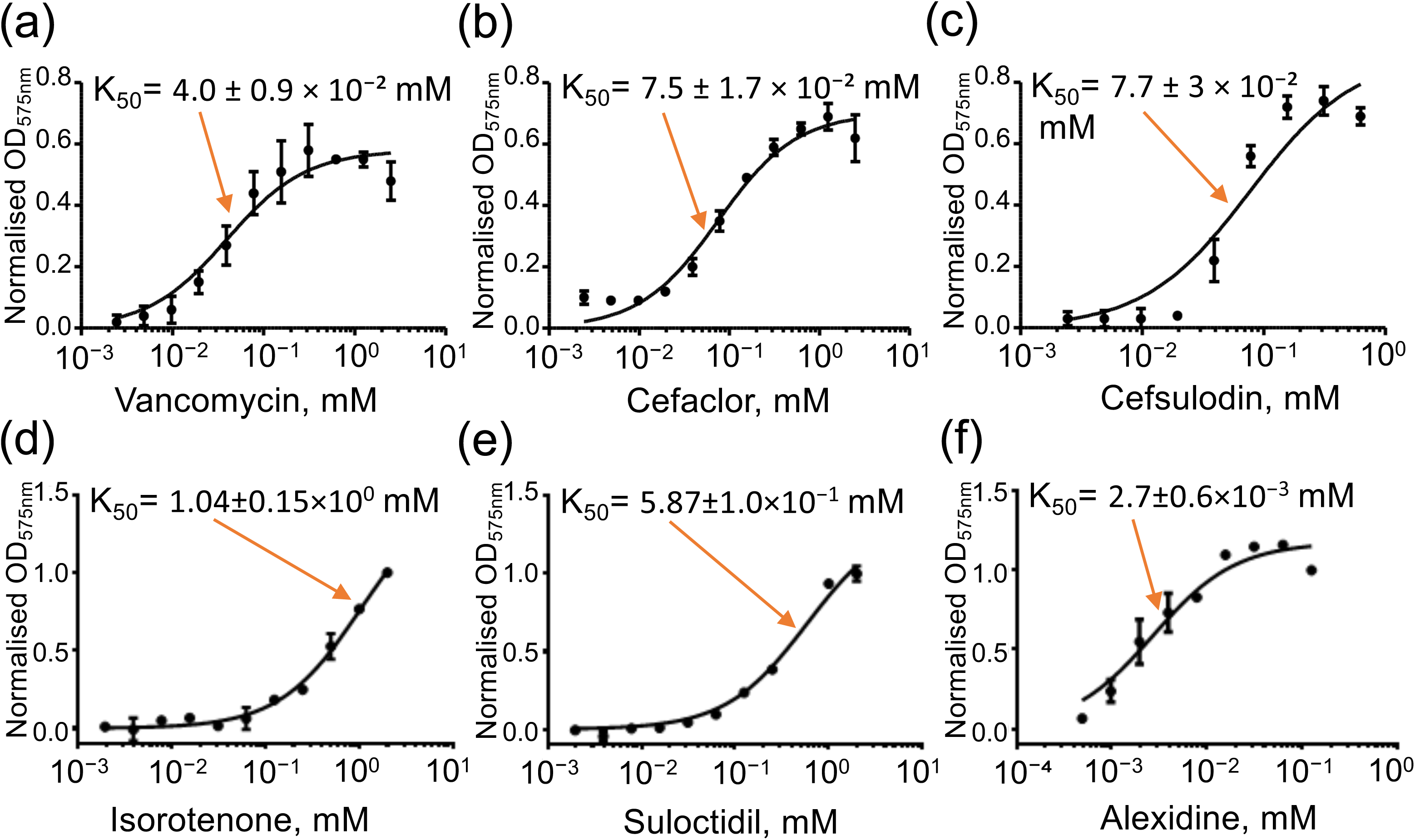
Permeabilisation curves determining the K_50_ for six compounds from the KD2-3840 library, identified in the primary screen. K_50_ marks the concentration at which OD_575_ absorbance rises exponentially, indicating the transition from low to high absorbance. The x-axis represents concentration (mM), and the y-axis denotes OD_575_nm absorbance of the CPR product across different concentrations. Vancomycin (a), cefaclor (b), cefsoludin (c), isorotenone (d), suloctidil (e), and alexidine (f) show a concentration-dependent positive readout in the CPRG assay. The sigmoidal curve exhibits a sharp increase, with K_50_ (arrow) marking the transition to high permeability. Among these, alexidine (f) reached K_50_ at the lowest concentration.

Among the non-antibiotic compounds, only alexidine exhibited concentration-dependent inhibition when titrated up to 0.025 mM (25 µM). Its permeabilisation constant (K_50_) was the lowest, at [(2.7 ± 0.6) × 10⁻³ mM (2.7 ± 0.6 µM)], among both non-antibiotic and antibiotic compounds (Figure 5). Concentration-response analysis of the six repurchased compounds confirmed that all active compounds permeabilised the bacterial cell wall in a concentration-dependent manner, with K_50_ values varying significantly (Table S1).

### Co-Permeabilisation Assay

Chloramphenicol, an inhibitor of bacterial protein synthesis that does not permeabilise the *E. coli* membrane, was tested to assess whether co-permeabilisation with screened compounds enhances its efficacy. The assay measured *β*-galactosidase activity via CPRG hydrolysis as a proxy for bacterial metabolism. The results reveal distinct interactions between chloramphenicol and the tested compounds, indicating potential synergies, antagonisms, or neutral effects that could inform combination therapy strategies. The assay demonstrated that chloramphenicol exhibited concentration-dependent effects on *E. coli* MG1655 when combined with 0.05 mM (50 µM) ampicillin, 0.002 mM (2 µM) alexidine, or 0.5 mM (500 µM) suloctidil. However, no activity was observed in the presence of 0.05 mM (50 µM) vancomycin, 0.5 mM (500 µM) isorotenone, or without a secondary agent (Figure 6). The results demonstrate that ampicillin, alexidine, and suloctidil enhance chloramphenicol’s efficacy through synergistic interactions, as shown by increased OD absorbance. In contrast, isorotenone, vancomycin, as demonstrated by the absence of a secondary agent, suggesting neutral or antagonistic interactions. These findings are essential for evaluating potential combination therapies and understanding how compounds can augment chloramphenicol’s effectiveness.

**Figure 6.**
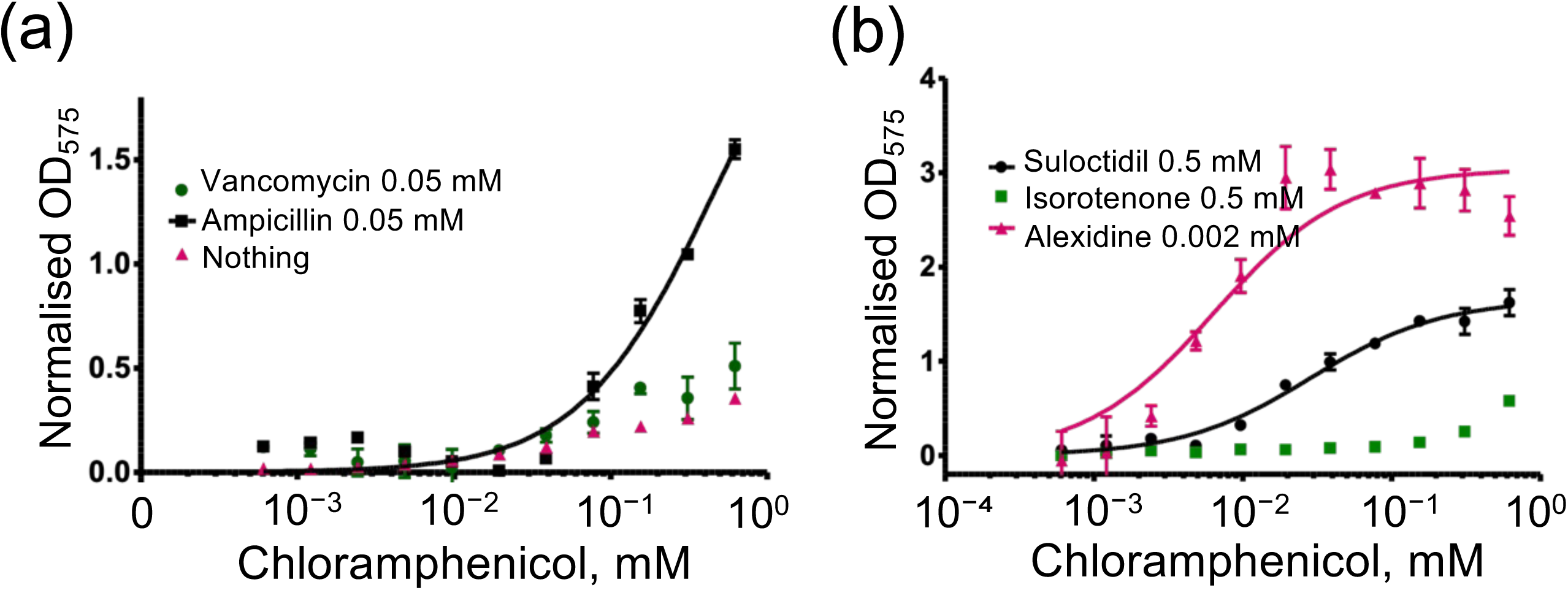
Effect of chloramphenicol on *E. coli* in the presence or absence of KD_2_ screen-identified compounds. (a) Co-permeabilisation assays assessed whether pre-treatment with selected compounds could enhance chloramphenicol activity, given its inability to permeabilise the *E. coli* membrane. Chloramphenicol did not exhibit a concentration-dependent effect when combined with 0.05 mM (50 µM) vancomycin, or without a secondary agent; however, Chloramphenicol showed a concentration-dependent effect when combined with 0.05 mM (50 µM) ampicillin. (b) Chloramphenicol exhibited a concentration-dependent effect when combined with 0.002 mM (2 µM) alexidine, or 0.5 mM (500 µM) suloctidil, while no enhancement was observed with 0.5 mM (500 µM) isorotenone. These findings highlight a potential strategy for evaluating combination therapies.

### The MIC and microscopic assays

The MIC assays confirmed the inhibitory effects of alexidine, suloctidil, and isorotenone, against *E. coli* and *P. aeruginosa*, with microscopic analysis revealing corresponding morphological alterations compared to untreated cells (Figure 7a, b). Alexidine exhibited the highest potency, with MIC values of 0.004 mM (4 µM) and 0.015 mM (15 µM) for *E. coli* and *P. aeruginosa*, respectively (Figure 7c). Cells treated with alexidine at MIC concentrations displayed a filamentous phenotype (Figure 7d, e), indicative of impaired cell division, often linked to envelope defects. Suloctidil demonstrated MIC values of 0.0125 mM (12.5 µM) and 0.5 mM (500 µM) for *E. coli* and *P. aeruginosa*, respectively (Figure 7f). Microscopic examination showed that suloctidil-treated cells became more spherical at sub-MIC concentrations (Figure 7g, h), with signs of lysis evident below the MIC, suggesting a potential defect in peptidoglycan integrity, crucial for osmotic stability. Isorotenone exhibited an MIC of 0.25 mM (250 µM) for both bacterial species (Figure 7i). Like suloctidil, isorotenone-treated cells adopted a more spherical morphology at sub-MIC concentrations (Figure 7j, k), with lysis occurring below the MIC. These findings further imply that the observed morphological changes may result from membrane perturbation, potentially compromising cell wall integrity and resistance to osmotic fluctuations, as elaborated in the discussion.

**Figure 7.**
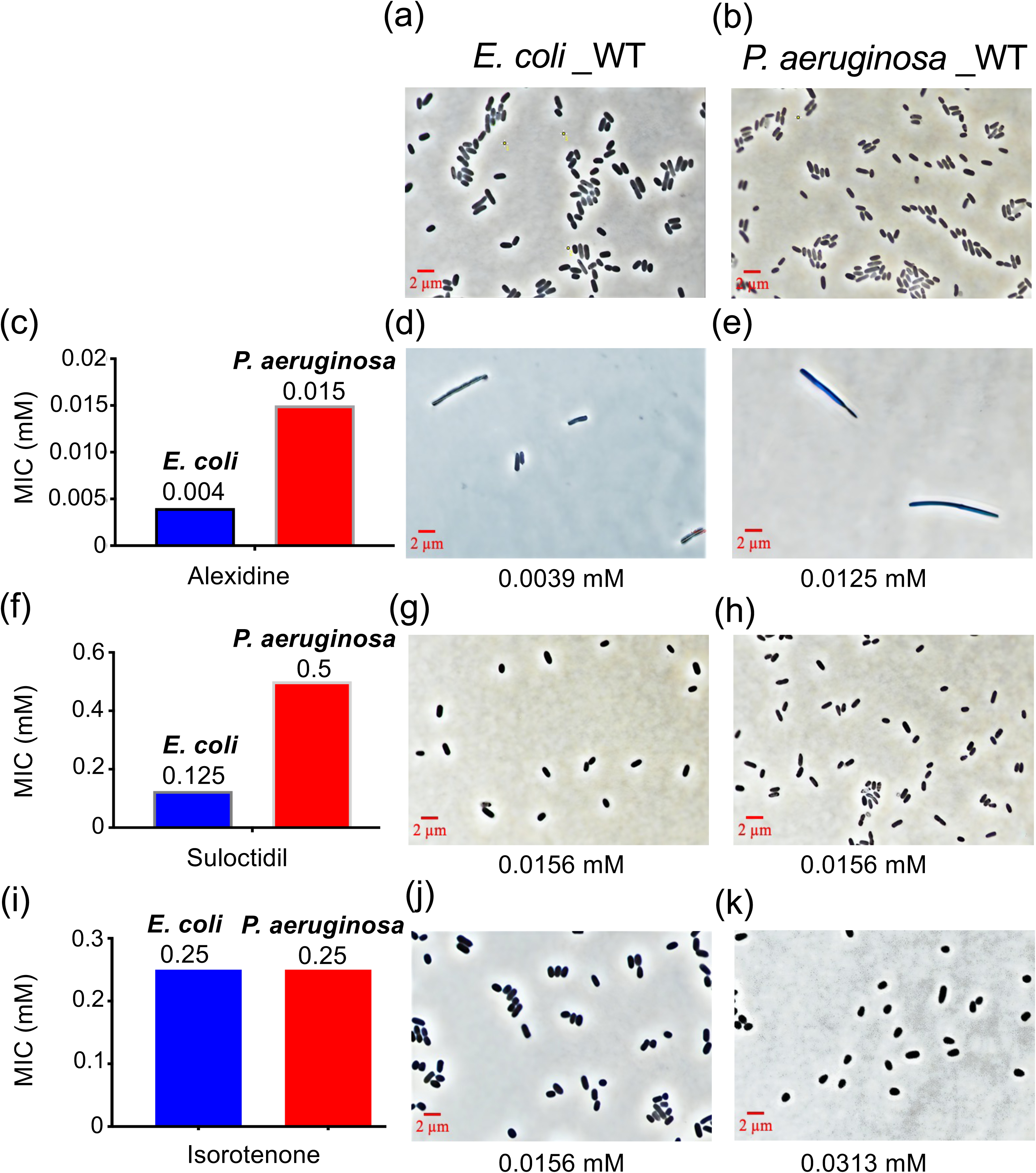
MIC and morphological effects of alexidine, suloctidil, and isorotenone in *E. coli* and *P. aeruginosa.* (a, b) Microscopy images showing the morphology of untreated bacterial cells. (c) Alexidine exhibited the lowest MICs: 0.004 mM (4 µM) for *E. coli* and 0.015 mM (15 µM) for *P. aeruginos*a. (d, e) Sub-MIC alexidine (0.0039, and 0.0125 mM) treatment induced filamentous morphology. (f) Suloctidil had MICs of 0.0125 mM (12.5 µM) for *E. coli* and 0.5 mM (500 µM) for *P. aeruginosa*. (g, h) Suloctidil-treated cells became more spherical and exhibited lysis below the MIC (0.0156 mM). (i) Isorotenone had an MIC of 0.25 mM (250 µM) for both species. (j, k) Isorotenone-treated cells also showed spherical morphology and lysis at sub-MIC (0.0156, and 0.0313 mM) concentrations. These results suggest membrane and cell wall disruption. The MIC values correspond to data obtained with the CPRG assay. Microscopy images taken with a 100x phase contrast objective.

## Discussion

This study introduces a validated, sensitive phenotypic assay in liquid media for the identification of candidate molecules targeting the GN bacterial envelope. The LacZ/CPRG assay, optimised for high-throughput screening (HTS), detects envelope permeability defects through OD at 575 nm. These findings provide a valuable tool for antibiotic discovery and may facilitate the identification of novel antibacterial agents.

Our assay enables high-performance screening of extensive compound libraries, exhibiting excellent Z-factor, S/B, and S/N ratios in LOPAC1280 and KD24761 screens. Optimised for 384- and suitable for 1536-well HTS^58^, it facilitates large-scale screening and identified 42 active compounds in the KD2^4761^ library (>25% inhibition), with 30 additional hits following overnight incubation. Notably, 90% of the hits were known antibiotics, with *β*-lactams comprising 80% (57 hits), 60% of which were unique. Screening of the KD2^3840^ library identified 57 active compounds (1.2% of the library), including repurchased antibiotics such as vancomycin, cefaclor, and sefsulodin, alongside non-antibiotic compounds such as alexidine, suloctidil, and isorotenone, confirmed concentration-dependent permeabilisation.

Among the identified compounds, alexidine emerged as the most potent, demonstrating significant envelope-disrupting activity at low concentrations. This bisbiguanide antiseptic destabilises bacterial membranes by binding to negatively charged phospholipids, enhancing membrane permeability, and inducing cell lysis, resulting in irregular cell morphology and bactericidal effects^59^. Our results suggest that alexidine is a promising candidate for targeting GN bacterial envelopes, with demonstrated antibacterial efficacy against *E. coli* and *P. aeruginosa*. Previously studied for its ability to neutralise lipopolysaccharides (LPS) and lipoteichoic acids (LTA), alexidine is also employed topically as a disinfectant against a broad spectrum of GN and GP bacteria, further supporting its potential as an antibacterial agent. Additionally, alexidine has exhibited strong antibacterial activity against both planktonic and biofilm forms of human fungal pathogens^59, 60^, as well as *Enterococcus faecalis* in dentin blocks^61^. However, its cytotoxicity has been assessed in various mammalian cell lines^60, 61^ and immune cells, such as primary bone marrow-derived macrophages^60^, where it exhibited slightly higher toxicity to macrophages (CC50 > 5 mg/L) compared to cell lines^60^. This underscores the need for further assessment of its selectivity and therapeutic index.

In addition to alexidine, suloctidil and isorotenone also permeabilised the bacterial envelope. Suloctidil, a peripheral vasodilator, disrupts lipid homeostasis and increases membrane fluidity, inducing bactericidal morphological changes^62^. Isorotenone, a rotenoid compound, inhibits respiration, depletes energy reserves, and potentially alters cell morphology, exhibiting antimicrobial, antifungal, and anticancer properties, although further investigation is warranted. These compounds induced distinct morphological changes at sub-minimum inhibitory concentrations (MICs), with suloctidil and isorotenone causing spherical forms and lysis, indicative of peptidoglycan layer disruption, while alexidine promoted a filamentous phenotype, suggesting a blockage in cell division. These morphological alterations suggest several potential mechanisms, including membrane disruption without peptidoglycan degradation, interference with cross-linking, or activation of autolysins. Moreover, they may involve effects on cytoskeletal elements, membrane–cell wall decoupling, or disruption of periplasmic homeostasis, leading to osmotic imbalances. Non-specific stress responses, such as membrane remodelling or division defects, may further contribute to these effects^63–65^. Further analyses, including peptidoglycan profiling, osmotic sensitivity assays, and electron microscopy, are needed to elucidate the primary mode of action.

Although many of the identified compounds are well-established antibiotics with limited potential for further development, the discovery of novel molecules such as alexidine, suloctidil, and isorotenone presents exciting opportunities. These compounds’ ability to destabilise the bacterial envelope offers potential for the development of new antibiotics or combination therapies. Our HTS platform, which facilitates rapid, large-scale screening, could play a pivotal role in identifying additional compounds targeting the bacterial envelope—a critical area for overcoming resistance mechanisms in Gram-negative pathogens^12, 66^. This study underscores the importance of developing novel strategies focused on the bacterial envelope, especially considering rising antimicrobial resistance^5^.

In comparison to other high-throughput antibacterial assays^29, 31, 32, 34, 37, 39, 41, 43^, and despite recent advancements in diagnostic tools for antibacterial resistance^18, 20, 21, 27, 28, 44–47^, our LacZ/CPRG assay stands out for its affordability, simplicity, speed, and high specificity in targeting the bacterial cell wall. Its superior sensitivity makes it invaluable for screening GN and potentially GP pathogens, with significant potential for integration into drug discovery pipelines. This assay could enhance the identification of novel antibiotics—whether by directly killing bacterial cells, blocking cell division, or augmenting the activity of other drugs via membrane permeabilisation. Its flexibility further allows the screening of larger compound libraries—from 384-well to 1536-well plates—thereby boosting throughput and expanding the discovery of new drug candidates.

In conclusion, our research demonstrates the efficacy of the LacZ/CPRG assay in identifying compounds that target the bacterial envelope, providing a powerful tool for antibiotic discovery. Consistent with the LacZ/ONPG assay, which was developed in a 96-well format to screen peptides as helper drug candidates that enhanced envelope permeability in *E. coli*^67^, our platform shows promise for identifying antibiotics suitable for both monotherapy and combination therapy. Amid the growing threat of antibiotic-resistant GN bacteria, this assay represents a critical step in developing effective therapeutics with a reduced risk of resistance evolution. Future efforts, in conjunction with deep learning methods^27, 28^, will expand screening to include a broader range of chemical libraries, advancing the development of envelope-targeting antibiotics to combat multidrug-resistant pathogens.

## Transparency declarations

None to declare.

## Supplementary data

Table S1 and Figures S1, S2, and S3 are available in the supplementary data at JAC Online.

## Acknowledgments

We would like to express our sincere gratitude to Dr. Jacques Thibodeau for his unwavering support during the preparation and publication of this manuscript. We also extend our thanks to the Department of Microbiology, Infectiology, and Immunology at the Université de Montréal, the Screening Division of CDRD, and the funding agencies NSERC and FRQS for their support.

## Supplementary data

The assay identified small molecules that targeted the envelope to kill cells or inhibit cell division, facilitating the development of bacteriolytic or bacteriostatic drugs, as illustrated in Figure S1. It also identified small molecules with permeabilisation activity, enabling the entry of other drugs into cells.

## Supplementary figure legends

**Figure S1.**
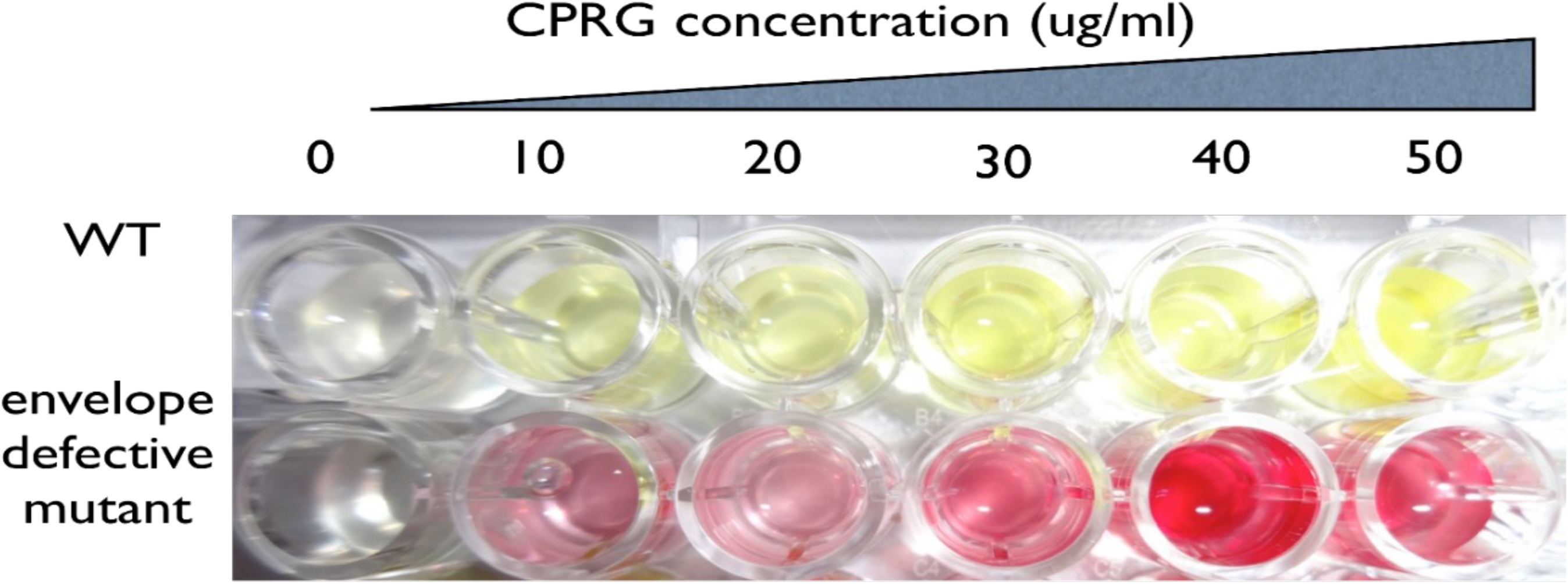
Optimising LacZ/CPRG assay for specificity and sensitivity in screening envelope-targeting antibiotics in LB media using 96-well plates. Microplate assay demonstrating efficient and specific detection of envelope permeability defects in envelope-defective E. coli mutant cells and wild-type (WT) cells. WT and envelope-defective E. coli mutant cells were cultured with 100 µM IPTG to induce LacZ production. Two hundred microliters of the bacterial cultures were transferred to microplate wells, followed by increasing concentrations of CPRG. The plate was incubated at room temperature for 30 minutes to allow colour development. The signal from both WT and mutant cells was concentration-dependent on CPRG and remained stable for over 3 hours. Results confirm that concentrations of 40 mg/L CPRG (chromogenic substrate) generate robust and reproducible signals.

**Figure S2.**
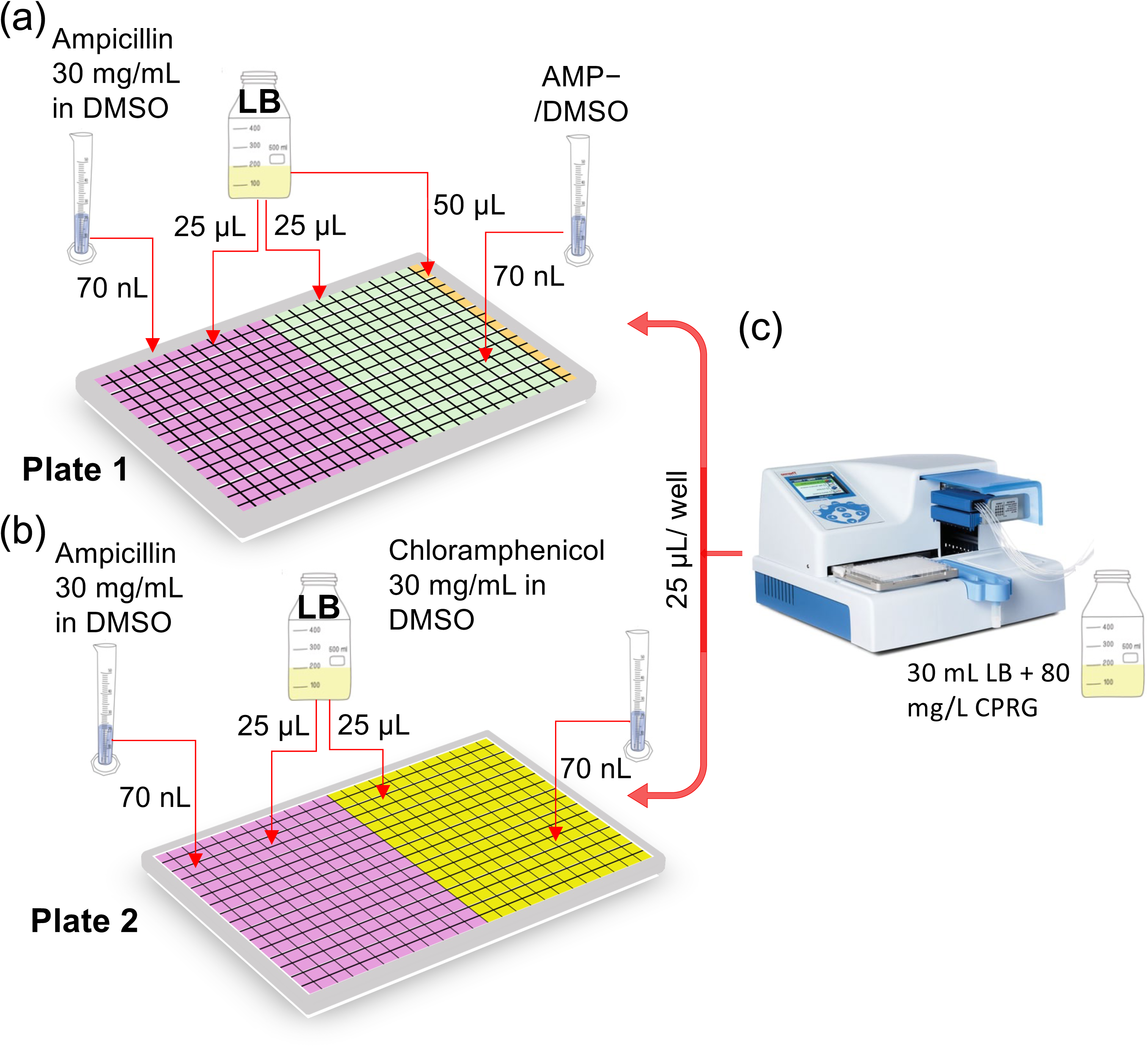
Plate preparation and compound transfer. Two 384-well plates were prepared with 25 µL of LB medium per well. (a) In Plate 1, wells received either 70 nL of ampicillin (30 mg/mL in DMSO) or 70 nL of DMSO (negative control, AMP−), with column 24 supplemented with 50 µL of fresh LB to monitor contamination. (b) In Plate 2, wells contained either 70 nL of ampicillin or chloramphenicol (30 mg/mL in DMSO). (c) A bacterial culture (30 mL) with CPRG (80 mg/L) was dispensed (25 µL per well) using a Thermo Combi dispenser, achieving a final CPRG concentration of 40 mg/L. Plates were incubated at room temperature with continuous agitation. Optical density at 575 nm (OD_575_) was measured after 3 hours (PowerWave XS, Biotek) and again after overnight incubation to assess cell wall leakage. Z’ values, signal-to-background (S/B), and signal-to-noise (S/N) ratios were calculated to evaluate assay quality.

**Figure S3.**
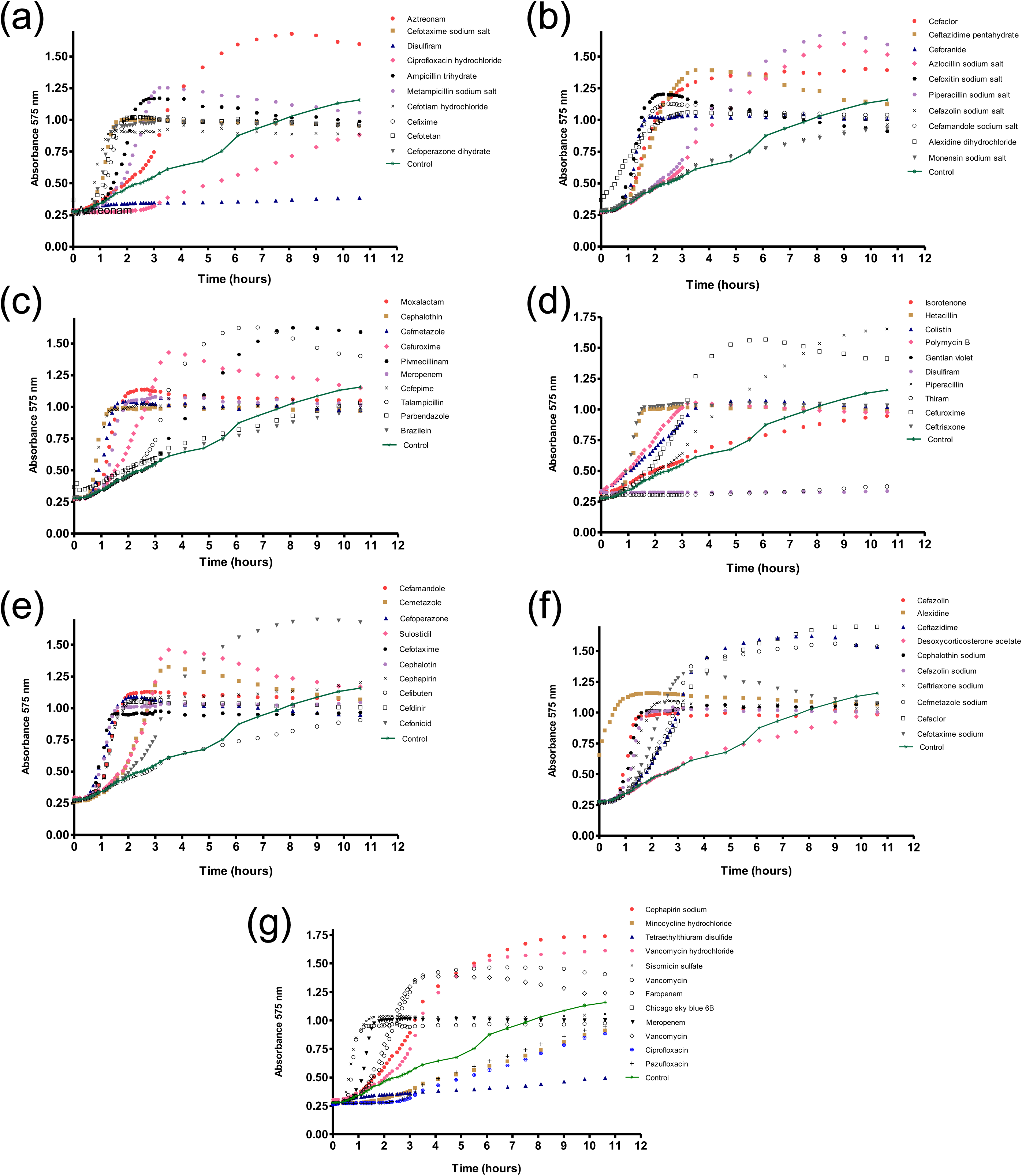
Progression of cell wall permeability (absorbance 575 nm) as a function of incubation time in the presence of compounds. All compound KD activities were organised into seven sets for clarity of display (a-g). The accumulation of background signal (DMSO only) is depicted as a solid green line. Absorbance was monitored at 575 nm for 11 hours for all active compounds to distinguish between those inducing slow leakage of the bacterial cell wall and those acting more rapidly. Each compound produced a distinct “leakage” curve. Twelve compounds were classified as “slow leakers”, nine as “moderate leakers”, and the remaining active compounds exhibited curves characteristic of fast-acting compounds.

## Supplementary Table

**Table S1.** Permeabilisation Constants (K50) of Six Selected Compounds.

| Compound name | Permeabilisation | Permeabilization |
| --- | --- | --- |
|  | constant, K<sub>50</sub> (mM) | constant, K<sub>50</sub> (μM) |
|  | Exponential Notation |  |
| Vancomycin | $(4.0 \pm 0.9) \times 10^{-2}$ | $40 \pm 9$ |
| Cefaclor | $(7.5 \pm 1.7) \times 10^{-2}$ | $75 \pm 17$ |
| Cefsoludin | $(7.7 \pm 3.0) \times 10^{-2}$ | $77 \pm 30$ |
| Isorotenone | $(1.04 \pm 0.15) \times 100^0$ | $1040 \pm 145$ |
| Suloctidil | $(5.87 \pm 1.00) \times 10^{-1}$ | $587 \pm 100$ |
| Alexidine | $(2.7 \pm 0.6) \times 10^{-3}$ | $2.7 \pm 0.6$ |

