## Supplemental Data 1 for "High-throughput LacZ/CPRG screen identifies novel potential antibiotics targeting Gram-negative bacterial envelopes to combat resistance"

**Running title: LacZ/CPRG high-throughput screen identifies new potential antibiotics**

Fardin Ghobakhlou[1*], Larbi Mokhtari[1], Tom A. Pfeifer[*2], and Catherine Paradis-Bleau[1]

[1]Department of Microbiology, Infectiology and Immunology, Faculty of Medicine, Université de Montreal, Quebec, Canada; [2]Biofactorial, Life Sciences Center, University of British Columbia, Vancouver, British Columbia, Canada.

*Corresponding authors:

The assay identified small molecules that targeted the envelope to kill cells or inhibit cell division, facilitating the development of bacteriolytic or bacteriostatic drugs, as illustrated in Figure S1. It also identified small molecules with permeabilisation activity, enabling the entry of other drugs into cells.

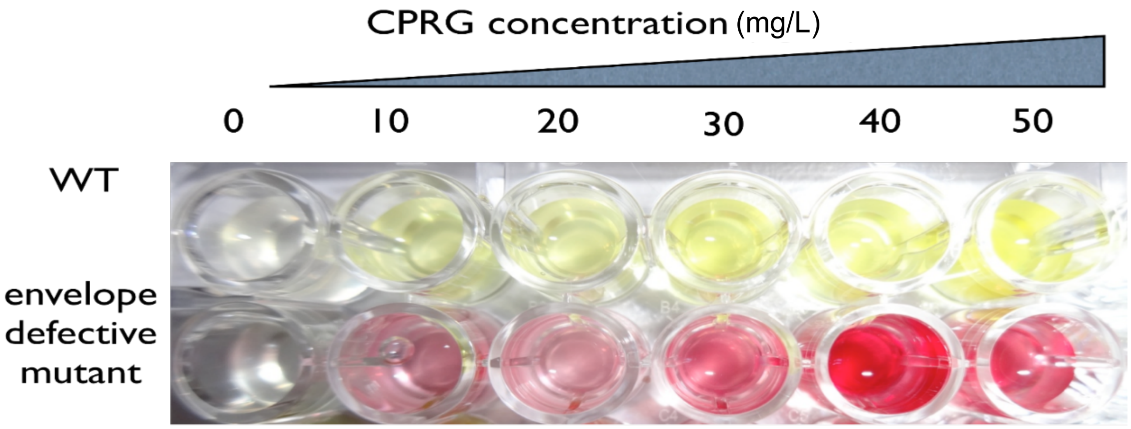


**Figure S1.** Optimising LacZ/CPRG assay for specificity and sensitivity in screening envelope-targeting antibiotics in LB media using 96-well plates. Microplate assay demonstrating efficient and specific detection of envelope permeability defects in envelope-defective *E. coli* mutant cells and wild-type (WT) cells. WT and envelope-defective *E. coli* mutant cells were cultured with 100 μM IPTG to induce LacZ production. Two hundred microliters of the bacterial cultures were transferred to microplate wells, followed by increasing concentrations of CPRG. The plate was incubated at room temperature for 30 minutes to allow colour development. The signal from both WT and mutant cells was concentration-dependent on CPRG and remained stable for over 3 hours. Results confirm that concentrations of 40 mg/L CPRG (chromogenic substrate) generate robust and reproducible signals.

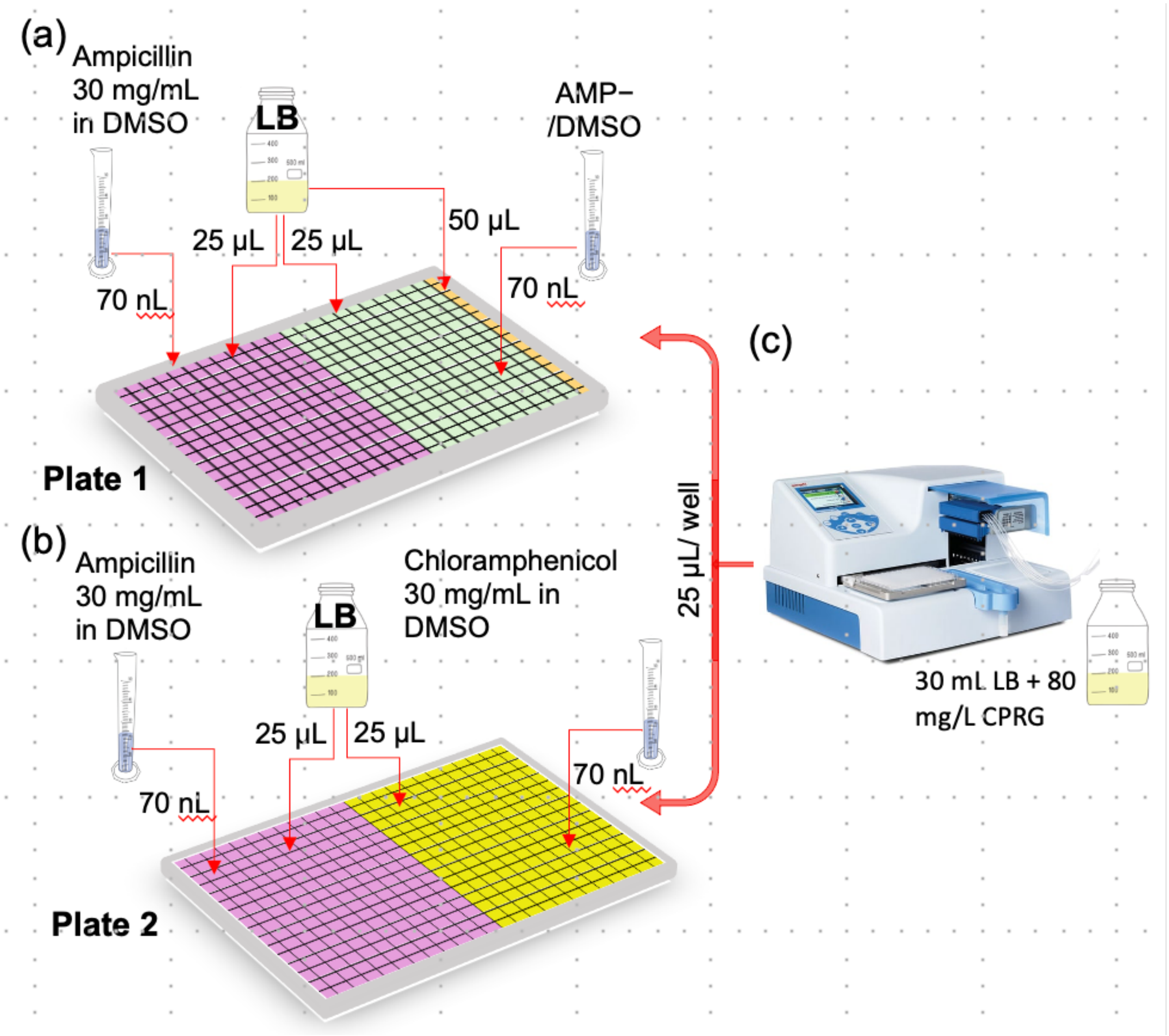


**Figure S2.** Plate preparation and compound transfer. Two 384-well plates were prepared with 25 µL of LB medium per well. (a) In Plate 1, wells received either 70 nL of ampicillin (30 mg/mL in DMSO) or 70 nL of DMSO (negative control, AMP−), with column 24 supplemented with 50 µL of fresh LB to monitor contamination. (b) In Plate 2, wells contained either 70 nL of ampicillin or chloramphenicol (30 mg/mL in DMSO). (c) A bacterial culture (30 mL) with CPRG (80 mg/L) was dispensed (25 µL per well) using a Thermo Combi dispenser, achieving a final CPRG concentration of 40 mg/L. Plates were incubated at room temperature with continuous agitation. Optical density at 575 nm ($OD_{575}$) was measured after 3 hours (PowerWave XS, Biotek) and again after overnight incubation to assess cell wall leakage. Z' values, signal-to-background (S/B), and signal-to-noise (S/N) ratios were calculated to evaluate assay quality.

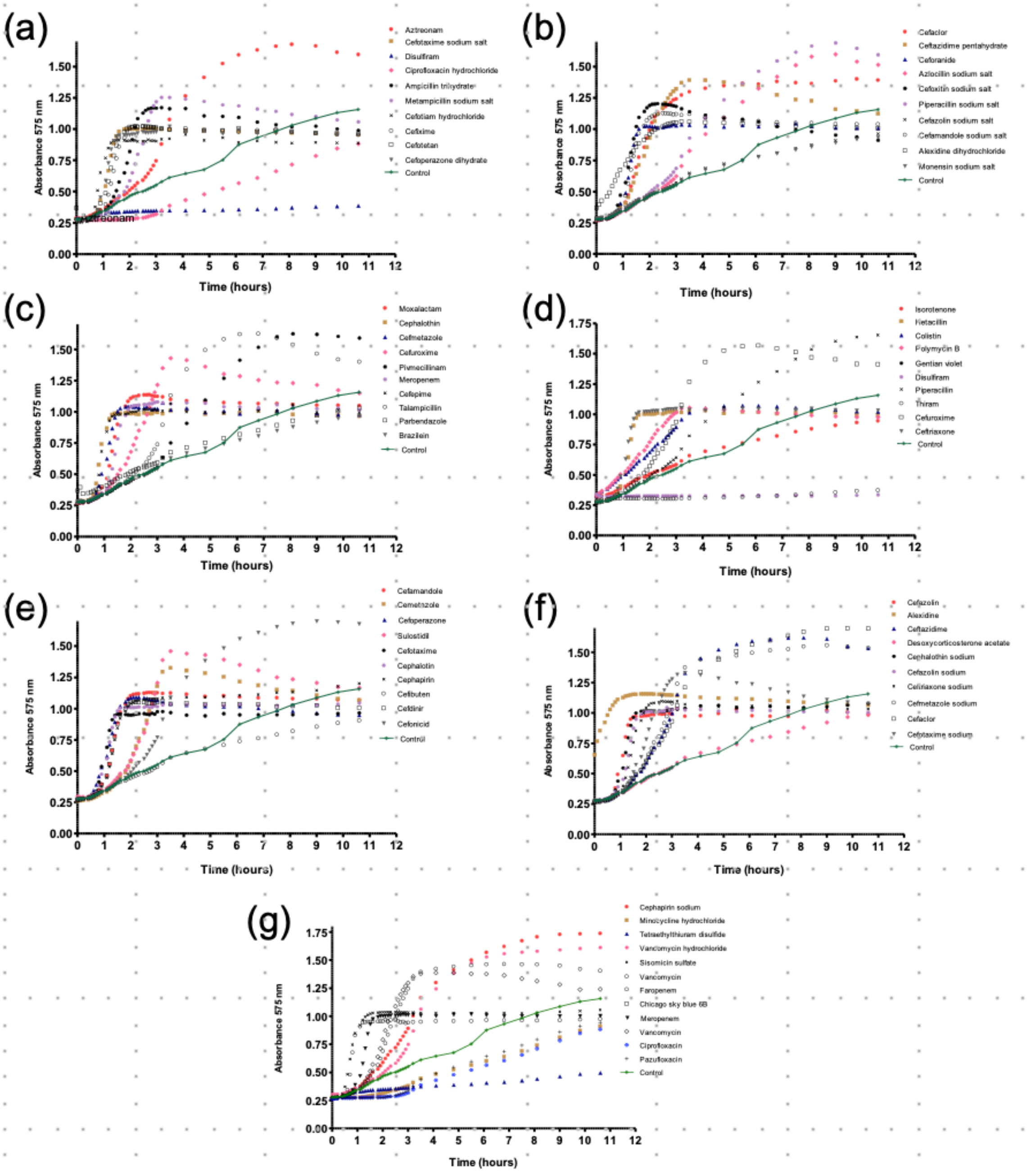


**Figure S3**. Progression of cell wall permeability (absorbance 575 nm) as a function of incubation time in the presence of compounds. All compound KD activities were organised into seven sets for clarity of display (a-g). The accumulation of background signal (DMSO only) is depicted as a solid green line. Absorbance was monitored at 575 nm for 11 hours for all active compounds to distinguish between those inducing slow leakage of the bacterial cell wall and those acting more rapidly. Each compound produced a distinct "leakage" curve. Twelve compounds were classified as "slow leakers", nine as "moderate leakers", and the remaining active compounds exhibited curves characteristic of fast-acting compounds.

**Table S1**. Permeabilisation Constants (K50) of Six Selected Compounds

| Compound name | Permeabilisation constant, $K_{50}$ (mM) Exponential Notation | Permeabilization constant, $K_{50}$ (μM) |
|---|---|---|
| Vancomycin | $(4.0 \pm 0.9) \times 10^{-2}$ | $40 \pm 9$ |
| Cefaclor | $(7.5 \pm 1.7) \times 10^{-2}$ | $75 \pm 17$ |
| Cefsoludin | $(7.7 \pm 3.0) \times 10^{-2}$ | $77 \pm 30$ |
| Isorotenone | $(1.04\pm0.15) \times 100^{0}$ | $1040 \pm 145$ |
| Suloctidil | $(5.87\pm1.00) \times 10^{-1}$ | $587 \pm 100$ |
| Alexidine | $(2.7\pm0.6) \times 10^{-3}$ | $2.7 \pm 0.6$ |
